# Resolving synaptic events using subsynaptically targeted GCaMP8 variants

**DOI:** 10.1101/2025.06.19.660577

**Authors:** Jiawen Chen, Junhao Lin, Kaikai He, Luyi Wang, Yifu Han, Chengjie Qiu, Jasmine M. Wheeler, Catherine M. Daly, Gregory T. Macleod, Dion Dickman

**Author notes:** Correspondence: Dion Dickman, Department of Neurobiology University of Southern California Los Angeles, CA 90089.

## Abstract

While genetically encoded Ca^2+^ indicators are valuable for visualizing neural activity, their speed and sensitivity have had limited performance when compared to chemical dyes and electrophysiology, particularly at synaptic compartments. We addressed these limitations by engineering a suite of next-generation GCaMP8-based indicators, targeted to presynaptic boutons, active zones, and postsynaptic compartments at the Drosophila neuromuscular junction. We first validated these sensors to be superior to previous versions and synthetic dyes. Next, we developed a Python-based analysis program, *CaFire*, which enables the automated quantification of evoked and spontaneous Ca²⁺ signals. Using *CaFire*, we show a ratiometric presynaptic GCaMP8m sensor accurately captures physiologically-relevant presynaptic Ca^2+^ changes with superior sensitivity and similar kinetics compared to chemical dyes. Moreover, we test the ability of an active zone-targeted, ratiometric GCaMP8m sensor to report differences in Ca²⁺ between release sites. Finally, a newly engineered postsynaptic GCaMP8m, positioned near glutamate receptors, detects quantal events with temporal and signal resolution comparable to electrophysiological recordings. These next generation indicators and analytical methods demonstrate that GCaMP8 sensors, targeted to synaptic compartments, can now achieve the speed and sensitivity necessary to resolve Ca^2+^ dynamics at levels previously only attainable with chemical dyes or electrophysiology.

## INTRODUCTION

Evolution’s ingenuity is evident in its exploitation of the calcium ion (Ca^2+^) as a potent secondary messenger, essential for controlling a diverse array of physiological events in every cell. The basis for Ca^2+^’s extraordinary signaling capacity lies in a formidable concentration gradient – maintained at nanomolar levels inside cells versus millimolar outside – that exceeds a thousand-fold. This carefully maintained difference allows for controlled Ca^2+^ entry to rapidly amplify cytosolic Ca^2+^ concentrations, in turn activating specific Ca^2+^-sensitive proteins to initiate vital downstream signaling pathways (Berridge et al., 2003, Clapham, 2007). In neurons, this Ca^2+^-driven control system achieves a pinnacle of sophistication. The powerful Ca^2+^ gradient is harnessed to orchestrate crucial neuronal processes, from shaping synaptic strength at dendritic spines and modulating gene expression in response to activity, to the precise triggering of neurotransmitter release via synaptic vesicle (Dittman et al., 2019, Jackman et al., 2017, McCarthy et al., 2024). Therefore, the ability to directly observe and interpret these dynamic Ca^2+^ signals in neurons provides a crucial lens, offering profound insights into the complex cellular operations that define neuronal function and flow of information through neural circuits.

Given the paramount importance of Ca^2+^ signaling in virtually all aspects of neuronal function, an extensive series of genetically-encoded Ca^2+^ indicators (GECIs) have been engineered with progressively enhanced speed and sensitivity. The archetypal GECI, GCaMP, ingeniously utilized a modified green fluorescence protein (GFP) flanked by the Ca^2+^ binding protein Calmodulin (CaM) and a Ca^2+^/CaM-binding M13-like peptide (Nakai et al., 2001). Upon Ca^2+^ binding, a conformational change in the CaM-peptide interaction allosterically modulates GFP fluorescence. While a conceptual breakthrough, early GCaMP iterations suffered from relatively slow kinetics, limited dynamic range, and modest fluorescence changes (Tian et al., 2009). Consequently, a variety of sophisticated protein engineering strategies were employed to bolster GCaMP performance, leading to significant improvements represented in the GCaMP6 and GCaMP7 series, which offered faster kinetics and higher signal-to-noise ratios (SNRs) (Dana et al., 2019, Chen et al., 2013). Finally, the subsequent development of the GCaMP8 series marked a substantial leap forward (Zhang et al., 2023). This enhanced performance was not merely incremental; it resulted from a deep understanding of the molecular mechanisms underlying GCaMP activation, involving meticulous optimization of the CaM/peptide interface for faster Ca^2+^ association and dissociation and other improvements that led to enhanced brightness and efficient conformational coupling (Zhang et al., 2024). These comprehensive engineering efforts yielded GCaMP8 variants that exhibit dramatically improved kinetics – both in rise and decay times – and sensitivity relative to previous generations. Indeed, their performance characteristics, including SNR, response speed, photostability, and linearity, became comparable or even superior to some of the best synthetic Ca^2+^ dyes, while retaining the crucial advantage of genetic targetability (Zhang et al., 2024). These innovations provide an opportunity to resolve Ca^2+^ dynamics at subcellular compartments with unprecedented resolution.

Many studies express GCaMP variants alone in the neuronal cytoplasm, a method that effectively captures the large, relatively slow Ca^2+^ fluxes in soma that correlate with action potential spiking (Akerboom et al., 2012, Peron et al., 2015, Ziv et al., 2013). However, Ca^2+^ transients within subcellular compartments, such as dendritic spines and presynaptic terminals, are characterized by significantly faster kinetics and occur within smaller, more restricted volumes than those in the general cytoplasm, posing distinct measurement challenges. Importantly, interpreting signals from GCaMPs expressed alone can be confounded by variations in sensor abundance due to uneven expression levels, differing protein stability, or photobleaching, any of which can obscure the accurate determination of true Ca^2+^ changes (McMahon et al., 2018). Ratiometric indicators are therefore an attractive alternative, as they typically involve co-expression of Ca^2+^-sensitive and Ca^2+^-insensitive fluorescent proteins in a stoichiometric manner (Zhang et al., 2021). By normalizing the Ca^2+^-dependent signal to the Ca^2+^-independent one, ratiometric measurements can control for these variations in sensor concentration and provide a more reliable quantification of Ca^2+^ levels and dynamics. Hence, an optimal approach for accurately capturing fast and local subsynaptic Ca^2+^ fluxes would target an advanced, ratiometric GCaMP sensor, engineered for high sensitivity and rapid kinetics, to the particular compartment being investigated.

The Drosophila neuromuscular junction (NMJ) is a powerful and genetically tractable model system for dissecting Ca^2+^ signaling dynamics at a glutamatergic synapse. Its utility is magnified by the ability to combine sophisticated genetic manipulation with GECIs, advanced imaging, and established electrophysiological approaches to monitor synaptic events with remarkable spatiotemporal resolution (Bellen et al., 2010, Frank et al., 2013). Early investigations into Ca^2+^ dynamics at the fly NMJ employed chemical dyes (e.g., OGB-1, Fura-2), which, despite their rapid kinetics, present challenges in controlling loading concentrations at synaptic compartments (Macleod et al., 2002). The advent of GECIs offered an opportunity for targeted and consistent expression at subcellular regions. Pioneering engineering of presynaptic GECIs enabled Ca^2+^ visualization at axonal boutons or active zones (Kiragasi et al., 2017, Cohn et al., 2015, Akbergenova et al., 2018), but were often non-ratiometric for Ca^2+^ sensing – complicating quantification – and employed older GCaMP variants with suboptimal kinetics and SNRs. Postsynaptic sensors evolved from using GCaMP2 to GCaMP6f, offering better sensitivity for detecting quantal events (Newman et al., 2017, Peled et al., 2014). More recently, GCaMP8f variants were employed (Perry et al., 2022, Li et al., 2021, Han et al., 2023), demonstrating promising sensitivity. However, other studies suggested that GCaMP8m may offer higher SNRs and similar kinetics, particularly in Drosophila (Zhang et al., 2023). These developments motivated us to engineer and validate optimal GECIs for the fly NMJ, aiming for performance that rivals chemical dyes for presynaptic imaging and electrophysiology for postsynaptic quantal event detection.

We have engineered and characterized a variety of next-generation GECI probes to resolve Ca^2+^ dynamics at pre- and post-synaptic compartments at the Drosophila NMJ. While an expected tradeoff between signal strength and kinetics was observed, GCaMP8m consistently delivered the highest responses with only modest reductions in speed relative to GCaMP8f. Notably, presynaptic GCaMP8m sensors captured physiologically important events with speed and sensitivity similar or superior to that of chemical dyes, while postsynaptic GCaMP8m probes achieved quantal event detection with similar resolution to electrophysiological recordings. These features establish targeted ratiometric GCaMP8m sensors and CaFire analysis as complementary tools for resolving synaptic events using purely optical approaches.

## RESULTS

### Engineering next generation GCaMP sensors targeted to synaptic compartments

To engineer the next generation of synaptically-targeted GCaMP sensors, we sought to improve upon previous sensors employed for Ca^2+^ imaging at the Drosophila NMJ. In particular, we focused on three cassettes. First, to target GCaMP to presynaptic boutons, GCaMP6s was fused to the synaptic vesicle protein Synaptotagmin (SYT), generating UAS-SYT::GCaMP6s (Cohn et al., 2015), schematized in **Fig. 1A**. More recently, this SYT::GCaMP fusion strategy was improved upon by replacing GCaMP6s with the faster and more sensitive GCaMP8f, and rendered ratiometric by fusion to the red-shifted, monomeric fluorescent protein mScarlet (Bindels et al., 2017), to generate UAS-<u>S</u>YT::mS<u>car</u>let::GCaMP<u>8f</u> (Scar8f) (Li et al., 2021) (**Fig. 1A**). To potentially improve this indicator, we replaced GCaMP8f with GCaMP8m, which is reported to exhibit higher sensitivity with only minor reductions in kinetics (Zhang et al., 2023), and replaced mScartlet1 with mScarlet3, which encode improvements in brightness and photophysical characteristics (Gadella et al., 2023), to make the ratiometric GECI Scar8m (**Fig. 1A**). Expression of each sensor with the motor neuron driver OK319-Gal4 and immunostaining demonstrated the expected trafficking to presynaptic boutons and co-localization with SYT (**Fig. 1B**).

**Figure 1:**
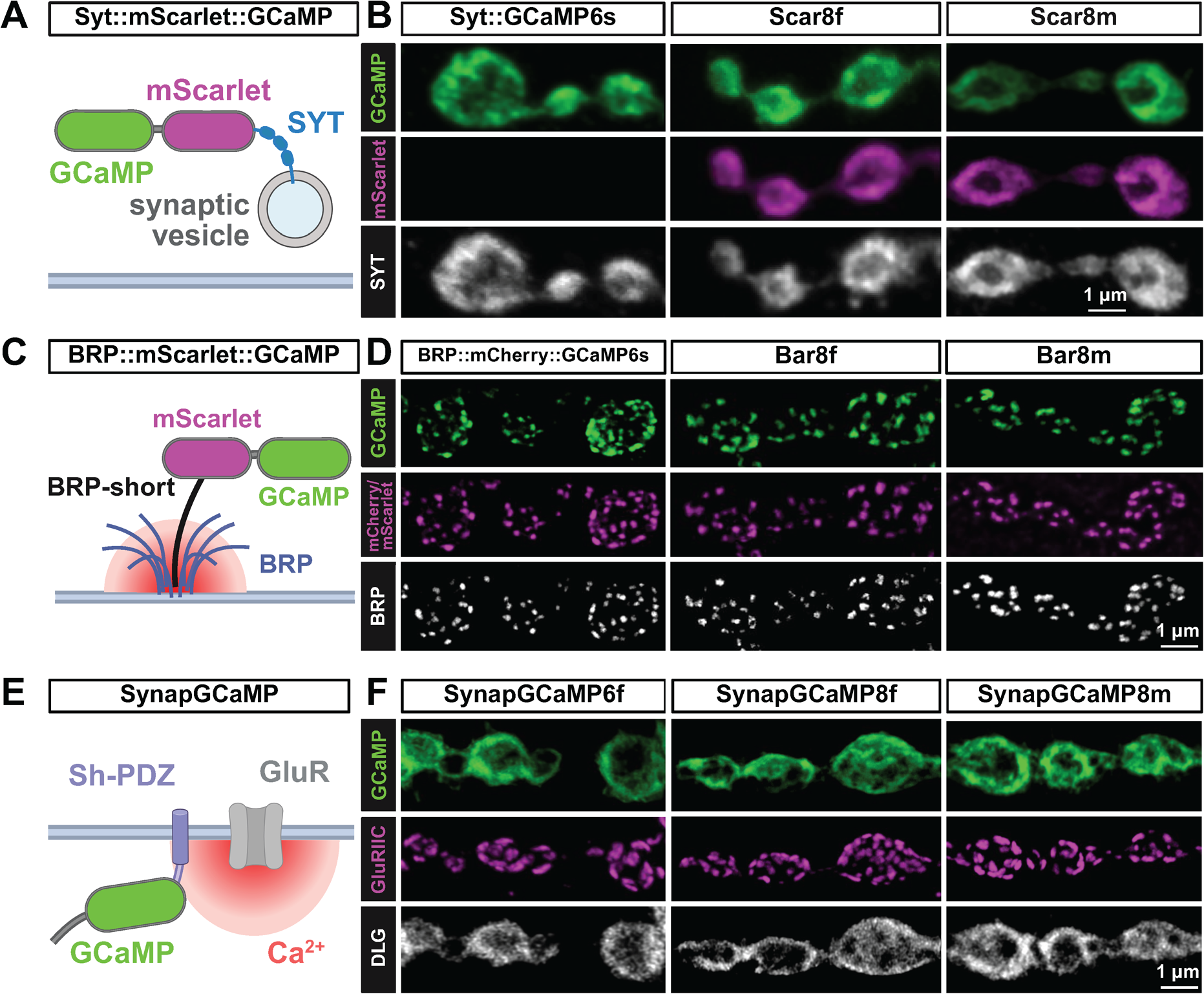
GCaMP indicators targeted to pre- and post-synaptic compartments. **(A)** Schematic of the presynaptic ratiometric Syt::mScarlet::GCaMP (Scar8f/Scar8m) Ca^2+^ indicators showing localization to synaptic vesicles via fusion to the Ca^2+^ sensor Synaptotagmin (SYT). **(B)** Representative images of NMJs expressing the indicated reporter driven in motor neurons with the OK319-GAL4 driver (*w^1118^*; *OK319-GAL4*/*UAS-Scar8f*) immunostained with anti-GFP (GCaMP) and anti-SYT. Note that endogenous mScarlet signals were obtained without antibody labeling. **(C)** Schematic of the BRP::mScarlet::GCaMP8f/8m (Bar8f/Bar8m) ratiometric Ca^2+^ indicator, which targets GCaMP to active zones via fusion to the BRP-short protein (Schmid et al., 2008). **(D)** Representative images of NMJs expressing the indicated reporter driven in motor neurons (*w*;*OK6-GAL4*/*Bar8f* and *w;OK6-GAL4/Bar8m*) immunostained with anti-GFP (GCaMP) and anti-BRP. Note that native mCherry or mScarlet signals were obtained without antibody labeling. **(E)** Schematic of the SynapGCaMP indicator, which targets GCaMP to postsynaptic compartments via a Shaker PDZ domain.(Newman et al., 2017) **(F)** Representative images of NMJs expressing the indicated reporter (*w*;*MHC-CD8-GCaMP6f-Sh/+*;*+*, *w*;;*MHC-CD8-GCaMP8f-Sh*/*+*, *w*;;*MHC-CD8-GCaMP8m-Sh*/*+*) immunostained with anti-GFP (GCaMP), -GluRIIC (glutamate receptors), and -DLG (postsynaptic density). All confocal images in this figure were deconvolved in Huygens (CMLE) prior to display (see methods).

Next, to target GCaMP to the sites of presynaptic Ca^2+^ influx, active zones (AZs), we improved upon a previous indicator, UAS-BRPshort::mCherry::GCaMP6s (Kiragasi et al., 2017). This strategy used a “short” fragment of the AZ scaffold Bruchpilot (BRP) (Schmid et al., 2008), which traffics to AZs, fused to mCherry and GCaMP6s for ratiometric Ca^2+^ imaging at AZs (**Fig. 1C**). To improve upon this design, we replaced mCherry with the improved mScarlet, and GCaMP6s with GCaMP8f to make UAS-<u>B</u>RPshort::mSc<u>ar</u>let::GCaMP<u>8f</u> (Bar8f) or GCaMP8m and mScarlet3 to make Bar8m (**Fig. 1C**). Expression of these sensors in motor neurons followed by immunostaining also confirmed the expected trafficking to AZs and co-localization with BRP (**Fig. 1D**). Finally, we worked to improve upon a postsynaptic Ca^2+^ sensor. A previous cassette called SynapGCaMP6f fused GCaMP6f to a Shaker PDZ motif to target the sensor near postsynaptic NMJ glutamate receptors (GluRs) under the control of a muscle-specific enhancer to enable “quantal imaging” of single synaptic vesicle release events (Newman et al., 2017) (**Fig. 1E**). A recent improvement replaced GCaMP6f with GCaMP8f (Han et al., 2022), and here we engineered GCaMP8m to make SynapGCaMP8m (**Fig. 1E**). Immunostaining of NMJs expressing each sensor confirmed proper trafficking and localization to postsynaptic densities near GluRs (**Fig. 1F**). In addition, GCaMP is a Ca^2+^ buffer and may therefore impact synaptic development and/or function (McMahon et al., 2018). Thus, we recorded from NMJs expressing the indicators introduced above and found that expression of the GCaMP sensors did not perturb synaptic transmission, with the exception of Scar8f, which showed a moderate reduction in EPSP amplitudes (**Fig. S1**). Collectively, these tools established the foundation of our study to go on to benchmark each sensor and determine the optimal indicator, and their speed and sensitivity, to resolve pre- and post-synaptic Ca^2+^ events and dynamics.

### Automated analysis of Ca^2+^ imaging data

Current approaches to analyze synaptic Ca^2+^ imaging data either repurpose software designed to analyze electrophysiological data or use custom software developed by groups for their own specific needs, including measuring spike frequency across many neurons (Müller et al., 2012, Giovannucci et al., 2019, Blum et al., 2021, Xing et al., 2018). We therefore developed a new Python-based software platform capable of automated detection and quantification of synaptic Ca^2+^ events. This program, which we named “CaFire”, analyzes imaging data followed by optional processing such as deconvolution (**Fig. 2A**). The Ca^2+^ imaging data are then imported into ImageJ to select Regions of Interest (ROIs), which are finally exported to a graphical program plotting event time and intensity data (e.g., Microsoft Excel; see methods for details). The CaFire software reads the Excel file and performs all downstream analysis.

**Figure 2:**
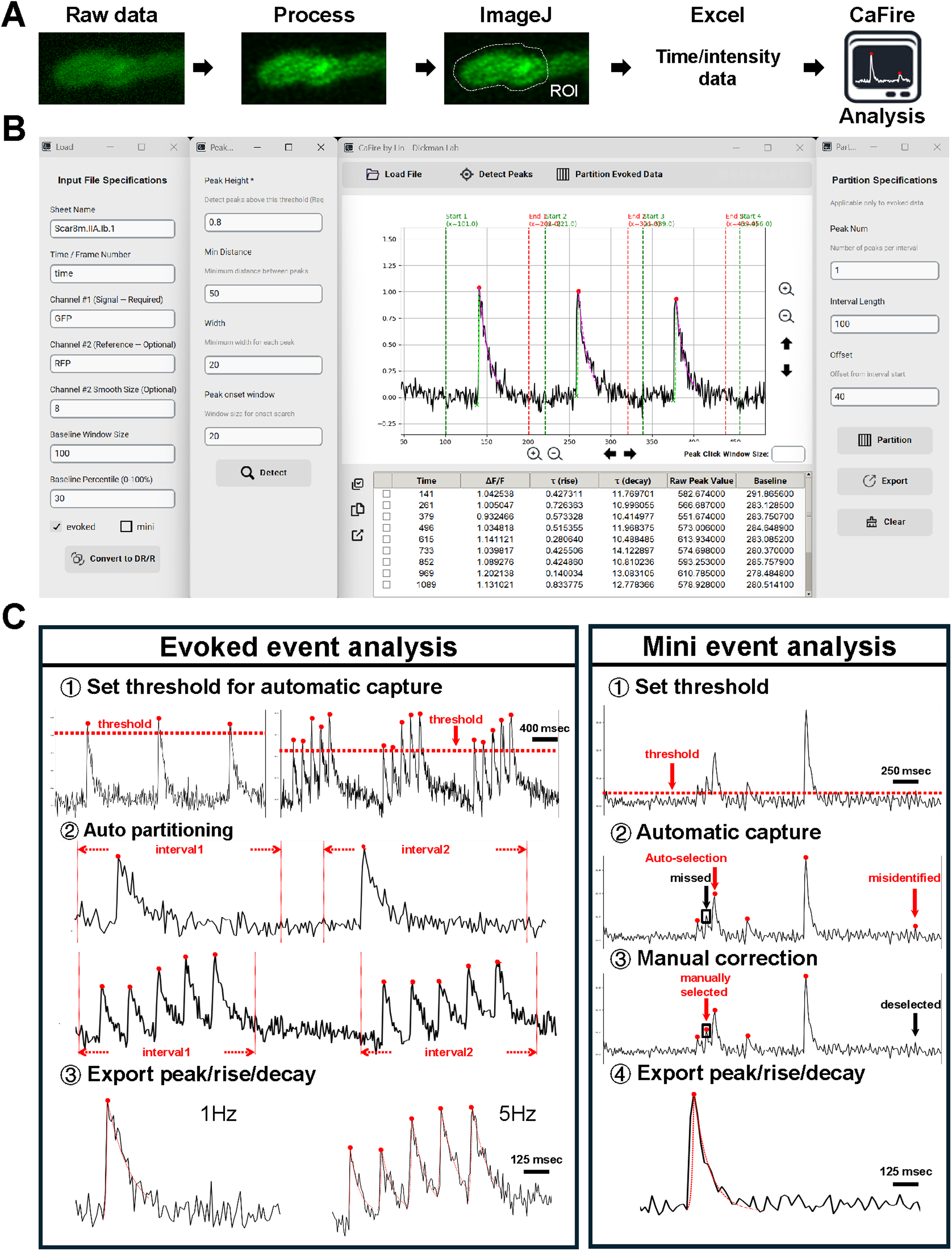
“CaFire” - a Python-based analysis program for quantifying synaptic Ca^2+^ imaging data. **(A)** Workflow showing how Ca^2+^ imaging data and downstream analysis is performed. Raw timelapse movies are processed with SVI Huygens software to correct and deconvolve image artifacts. ROIs are then selected in ImageJ Fiji software and intensities are extracted. Data are exported to Excel for manual inspection or directly analyzed using CaFire software. **(B)** Screenshots of the CaFire user interface. The left panels allow users to specify input file properties and analysis settings, and the second panel from the left shows the peak-detection criteria. The central panel displays fluorescence traces with automatically detected Ca²⁺ events overlaid and the event-based partition results. Detected events and their extracted parameters are listed in the data table below. The right panel shows the partition specifications used to segment evoked responses. **(C)** Examples of two distinct analysis pipelines implemented in CaFire. *Evoked event analysis*: (1) Thresholds are set for automatic peak detection; (2) Events are automatically partitioned based on stimulus intervals (e.g., 1 Hz and 5 Hz); (3) Parameters such as peak amplitude, rise time constant (τ_rise_), and decay time constant (τ_decay_) are calculated using exponential fits. *Mini event analysis*: (1) Users define amplitude thresholds for event detection; (2) CaFire automatically identifies candidate events; (3) Missed or misidentified events can be manually corrected; (4) Event parameters are exported for each validated event.

CaFire can analyze both presynaptic evoked and postsynaptic quantal Ca^2+^ events with a user-friendly graphical interface that enables parameter adjustment and event visualization (**Fig. 2B**). For both quantal and evoked events, peak detection is based on customizable parameters, including threshold amplitude, peak width, and minimum inter-peak distance. These parameters can be adjusted to optimize detection sensitivity and specificity, while misidentified events can be manually corrected. Once events are identified, CaFire then quantifies key parameters, such as peak amplitude (ΔF/F), rise time constants based on an exponential growth formula, and decay time constants using a natural logarithmic decay formula (**Fig. 2C**; see methods). In addition, evoked Ca^2+^ events can be partitioned at different stimulation frequencies (e.g., 1 Hz, 5 Hz) to assess frequency-dependent dynamics. All quantification results are displayed in real-time and can be exported in a tabular format for downstream statistical analysis. This automated workflow significantly reduces analysis time and user variability in Ca^2+^ imaging data analysis when compared to a variety of other platforms we have tried, providing uniform, accurate, and reproducible measurements. CaFire’s ability to analyze both evoked and quantal events makes it a versatile tool for quantifying synaptic Ca^2+^ imaging data.

### Benchmarking presynaptic Ca^2+^ sensors

We first sought to directly compare the performance of Scar8f with chemical Ca^2+^ indicator Oregon Green BAPTA-1 (OGB-1) under the same conditions. Images of the type-Ib terminal on muscle #6 (MN6/7-Ib) terminals were captured at 303 frames-per-second on a confocal microscope (**Fig. 3A**), evoking fluorescence transients at 1 Hz. In all aspects, Scar8f matched or exceeded the performance of OGB-1 (**Fig. 3B-E**). Specifically, GCaMP8f displayed higher sensitivity compared to OGB-1 and exhibited a faster rate of decay (**Fig. 3B,C**). The terminals were also challenged with a 21.3 Hz train of stimuli, where individual Ca^2+^ transients were easily discerned in both GCaMP8f and OGB-1 traces (**Fig. 3D**). However, GCaMP8f fluorescence summated to a far greater degree, representing greater frequency facilitation of the Ca^2+^ transients (**Fig. 3D,E**). Finally, we compared the evoked Ca^2+^ signals from cytosolic GCaMP8m (RSET-8m, (Reagents et al., 2024)) to Scar8m (**Fig. 3F**). We observed an almost two-fold increase in the amplitude of the Scar8m response in Scar8m compared to cytosolic GCaMP8m (**Fig. 3G,H**). Thus, Scar8f/m, targeted to synaptic vesicles, exhibits superior sensitivity and kinetics compared to the chemical dye OGB-1 and cytosolic GCaMP8m indicators.

**Figure 3:**
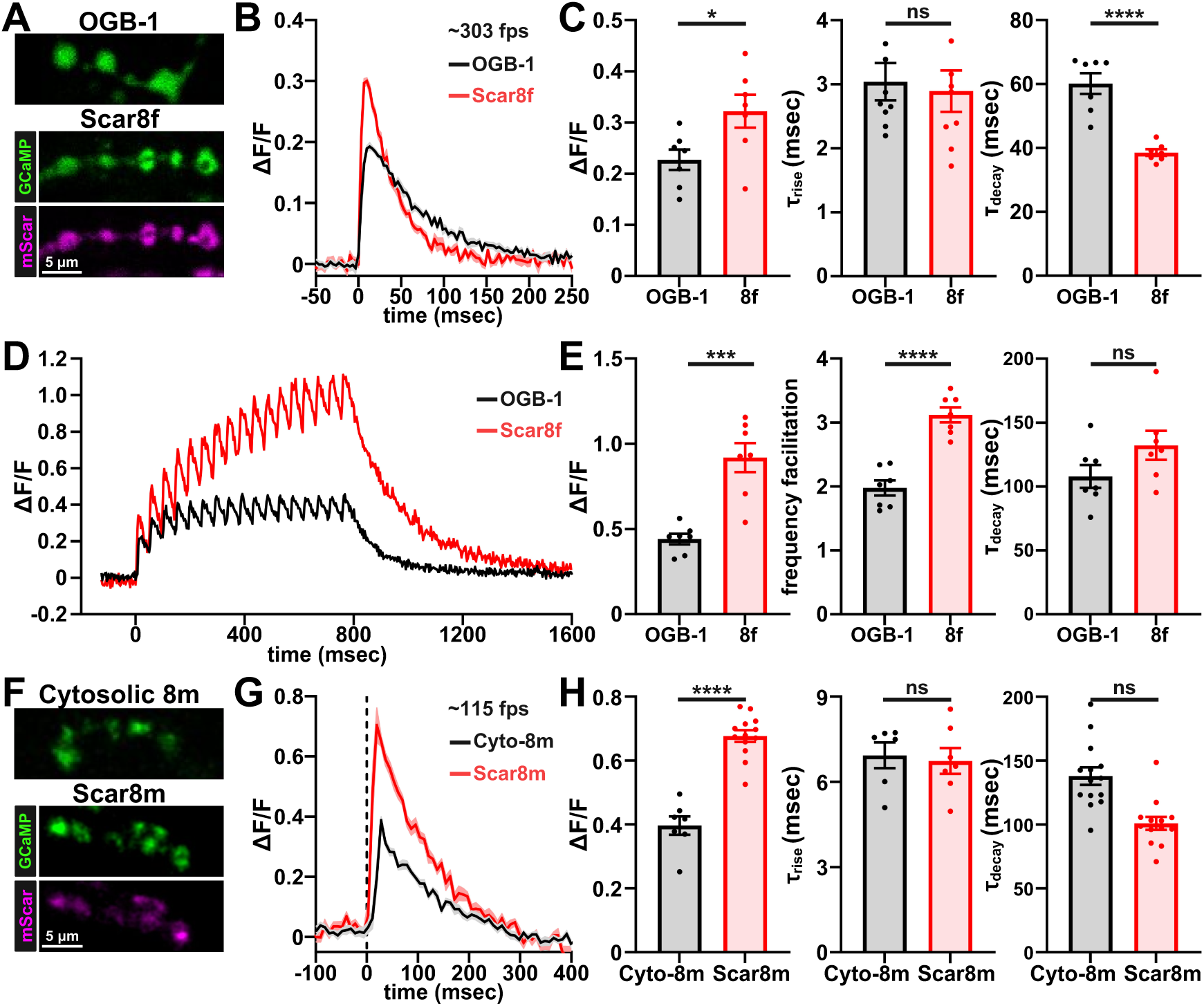
Benchmarking presynaptic GCaMP8 to synthetic dyes and cytosolic sensors. **(A)** Confocal images showing presynaptic loading of the synthetic dye OGB-1 and presynaptic expression of Scar8f at MN-Ib boutons. **(B)** Example single-AP evoked presynaptic Ca²⁺ transients reported by OGB-1 (black) and GCaMP8f (red). Traces represent the average of 30 APs from a single preparation acquired at 303 fps. **(C)** Quantification of response amplitude (ΔF/F), rise time, and decay times from single-AP Ca²⁺ transients reported by OGB-1 and Scar8f. **(D)** Example train-evoked Ca²⁺ transients reported by OGB-1 (black) and Scar8f (red) during three trains of 10 APs delivered at 21.3 Hz. **(E)** Quantification of train-evoked Ca²⁺ signals showing amplitude (ΔF/F) of the final Ca²⁺ transient in the train (17th), measured from baseline; frequency facilitation, calculated as the amplitude of the 17th transient divided by the amplitude of the 1st; and decay time constant (τ_decay_) measured after the final transient in the train. **(F)** Confocal NMJs images showing presynaptic expression of the cytosolic sensor RSET-8m (Cytosolic GCaMP8m) and Scar8m. **(G)** Example single-AP evoked Ca²⁺ transients reported by RSET-8m (black) and Scar8m (red) acquired at ∼115 fps. **(H)** Quantification of amplitude (ΔF/F), rise time (τ_rise_) and decay time constants (τ_decay_) from single-AP Ca²⁺ transients reported by RSET-8m (*w;OK319-GAL4/+; UAS-RSET-GCaMP8m/+*) and Scar8m (*w;OK319-GAL4/+; UAS-Syt::mScarlet::GCaMP8m/+*). Shaded traces and bars show mean ±SEM. Statistical comparisons are unpaired two-tailed t-tests; significance is indicated as *p<0.05, **p<0.01, ***p<0.001, ****p<0.0001; ns, not significant. Detailed statistics including p-values are provided in Table S1.

Next, we systematically evaluated the performance of the presynaptically targeted Ca^2+^ indicators SYT::GCaMP6s, Scar8f, and Scar8m at MN-Ib terminals. Live confocal imaging confirmed robust expression of all sensors at presynaptic boutons, with colocalized GCaMP and mScarlet fluorescence (**Fig. 4A,B**). Ca^2+^ transients were elicited by single action potential (AP) electrical stimulation, and fluorescence responses were recorded and analyzed (**Fig. 4C,D**). Scar8m produced the highest amplitudes of GCaMP/mScarlet (ΔR/R) ratios, with GCaMP8m showing 345.7% higher SNR over GCaMP6s, and 55.7% higher compared to GCaMP8f (**Fig. 4D,E**). Both Scar8f and Scar8m clearly outperformed GCaMP6s in both sensitivity and speed (**Fig. 4D,E**), as expected. Importantly, while GCaMP8f exhibited slightly faster decay time constants (66.6 msec vs 99.2 msec), rise time constants were not significantly different between the two sensors (5.3 msec vs 7.0 msec) (**Fig. 4E**). Thus, Scar8m exhibited an optimal balance between kinetics and signal strength, depending on the objectives of the experimental approach.

**Figure 4:**
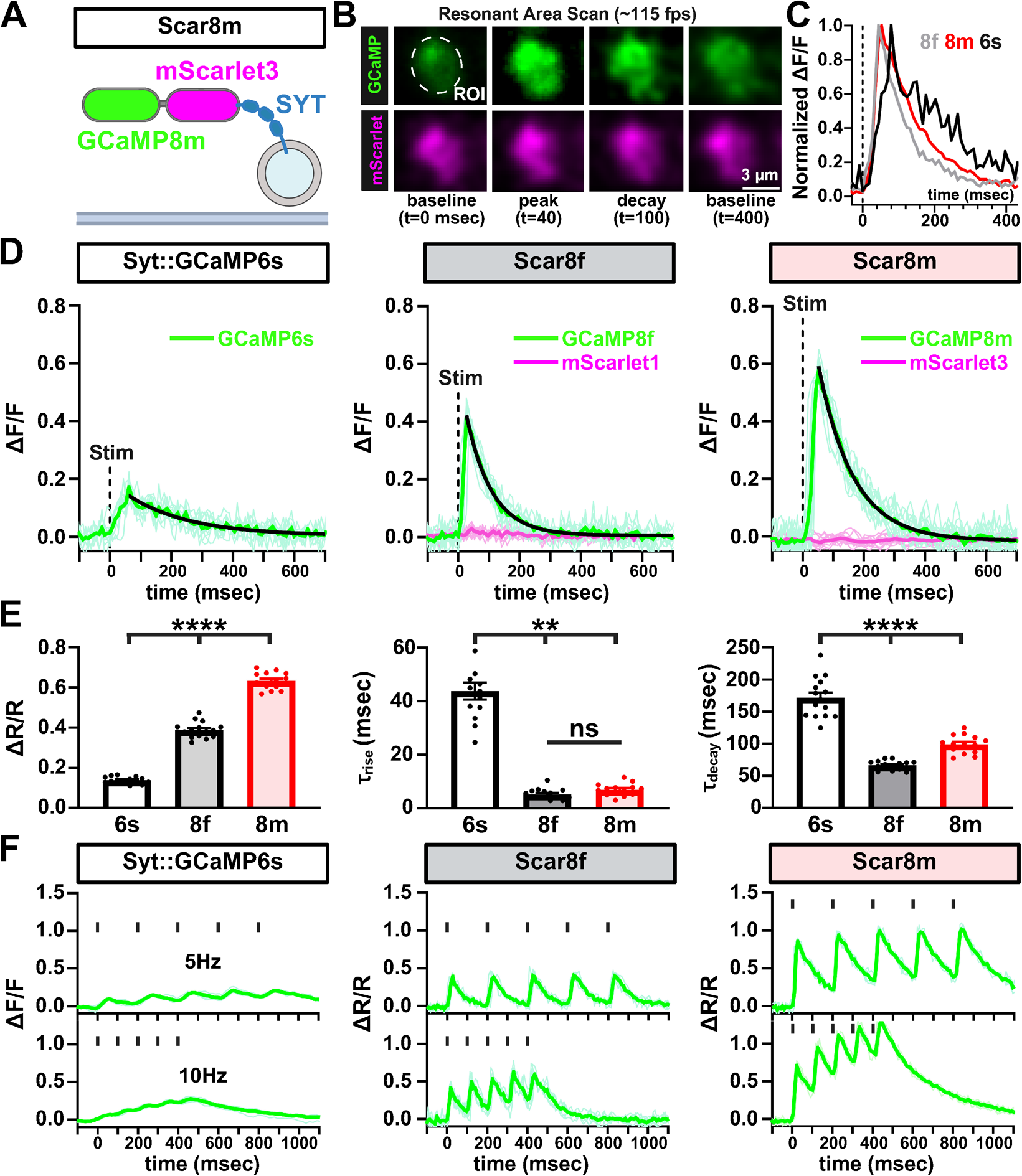
Scar8m is an optimal presynaptic Ca^2+^ indicator. **(A)** Schematic of the UAS-Syt::mScarlet3::GCaMP8m (Scar8m) ratiometric Ca²⁺ indicator consisting of mScarlet3 and GCaMP8m targeted to synaptic vesicles via fusion to Synaptotagmin (SYT). **(B)** Representative images of a MN-Ib bouton expressing Scar8m (*w*;*OK319-GAL4*/*+*;*Scar8m*/*+*) resonant scanned at ∼115 fps. Fluorescence from GCaMP8m (green) and mScarlet3 (magenta) is shown at baseline, peak, decay, and recovery to baseline. **(C)** Normalized Ca²⁺ signals following single action potential (AP) stimulation. Averaged traces compare the kinetics of GCaMP6s, GCaMP8f, and GCaMP8m. **(D)** Representative GCaMP and mScarlet signals recorded from Syt::GCaMP6s (GCaMP only), Scar8f, and Scar8m (GCaMP plus mScarlet) in response to single AP stimuli. Thin traces are sequential single-trial sweeps from the same NMJ; the thick colored trace is the mean trace after stimulus alignment. The black curve indicates a single-exponential decay fit used to estimate τ. **(E)** Quantification of average peak amplitude (ΔR/R, GCaMP/mScarlet ratios), rise time constant (τ_rise_), and decay time constant (τ_decay_) from the indicated sensors. Scar8m yields significantly higher peak ΔR/R signals, similar rise time kinetics, and a modestly slower decay compared to Scar8f. **(F)** Presynaptic Ca²⁺ responses of the indicated sensors to 5 Hz and 10 Hz stimulation trains. Vertical black ticks above the traces indicate the timing of stimulation pulses. Error bars represent ±SEM. Significance is indicated as *p<0.05, **p<0.01, ***p<0.001, ****p<0.0001; ns, not significant; detailed statistics including p-values are presented in Table S1.

To assess sensor performance during repetitive activity, we tested responses to 5 stimuli at 5 Hz and 10 Hz stimulation trains. While responses were muted using SYT::GCaMP6s, Scar8f and Scar8m consistently demonstrated robust responses and separation across repetitive stimulation (**Fig. 4F**). Together, we conclude that because of the superior sensitivity, ratiometric properties, and fast kinetics, Scar8m is an optimal GECI to quantify stimulated Ca^2+^ responses from presynaptic terminals.

To determine whether the sensitivity of Scar8m is sufficient to quantitatively report biologically relevant differences in evoked presynaptic Ca^2+^ responses, we compared evoked responses between MN-Ib and MN-Is terminals. Previous studies using chemical indicators have shown MN-Is exhibits ∼2-3x increased Ca^2+^ responses over MN-Ib (Lu et al., 2016, He et al., 2023). First, we confirmed robust Scar8m expression in both MN-Ib and -Is using the OK319-Gal4 driver (**Fig. 5A,B**). Next, Scar8m responses to single AP stimulation revealed a >two-fold increase at MN-Is terminals compared to MN-Ib (**Fig. 5C,D**). Next, we assessed changes in presynaptic Ca^2+^ responses at MN-Ib in wild type and *GluRIIA* mutants, in which a process called presynaptic homeostatic potentiation (PHP) is known to be induced (He et al., 2025, Davis et al., 2015). In PHP, diminished postsynaptic GluR functionality is offset through a homeostatic signaling system which enhances neurotransmitter release (Goel et al., 2021, Delvendahl et al., 2019). PHP induces an increase in presynaptic Ca^2+^ influx to promote additional synaptic vesicle release (Müller et al., 2012, Chien et al., 2025), where chemical indicators such as OGB-1 have shown ∼30% increase Ca^2+^ levels after PHP (Müller et al., 2012). Importantly, Scar8m responses in *GluRIIA* mutant MN-Ib boutons showed significantly elevated ΔR/R amplitudes compared to wild type, with a ∼53% enhancement (**Fig. 5E,F**). Thus, Scar8m can resolve biologically relevant differences in presynaptic Ca^2+^ with high sensitivity, on par or exceeding what chemical indicators have shown.

**Figure 5:**
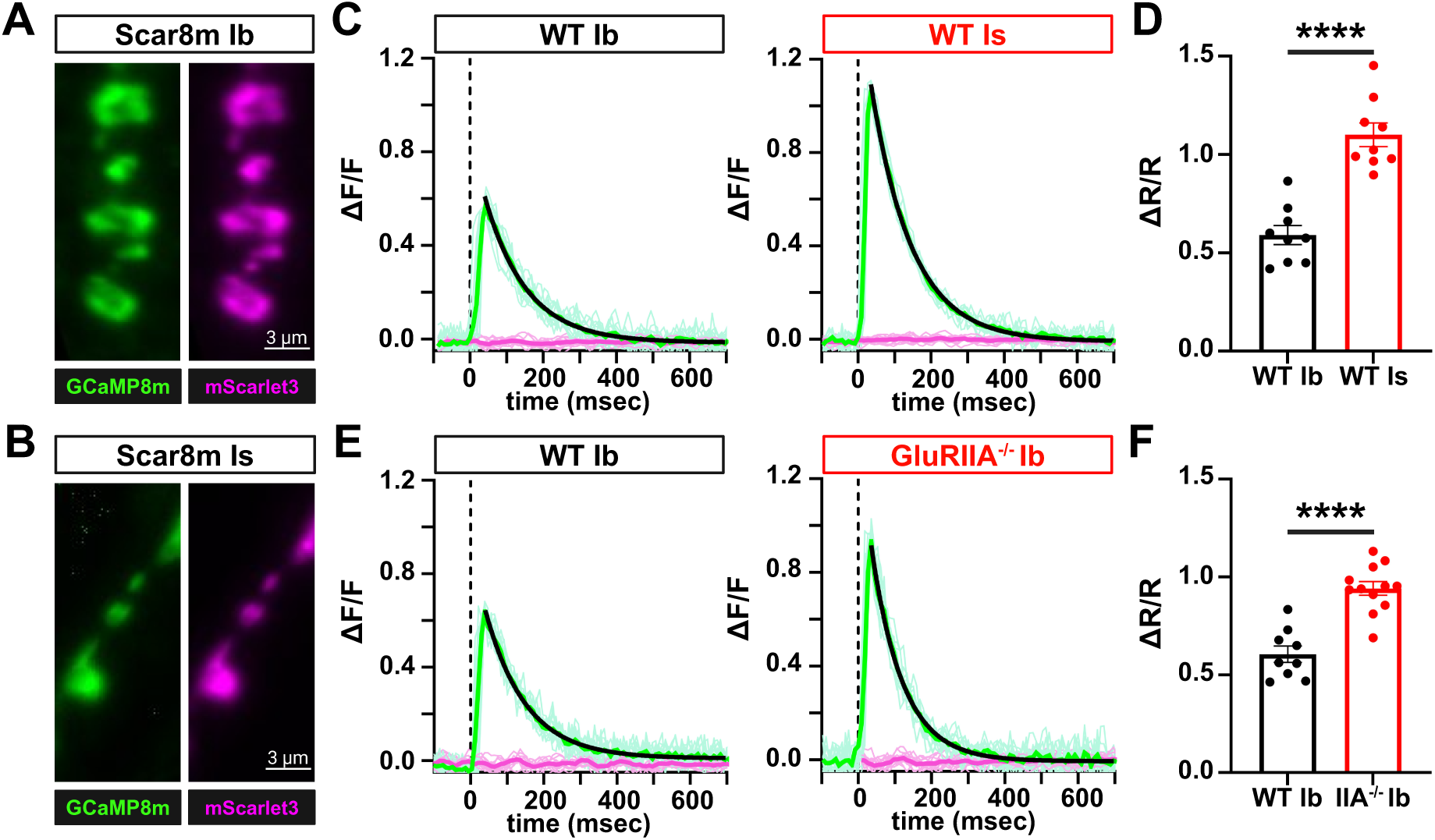
Scar8m captures differences in Ca^2+^ levels between motor neuron subtypes and after plasticity. (A,B) Representative images of Scar8m expressed at both MN-Ib (A) and MN-Is (B) motor neuron subtypes immunostained with anti-GFP. **(C)** ΔF/F traces of GCaMP8m and mScarlet3 responses from single AP stimulation at MN-Ib and MN-Is, with ∼2x higher responses observed at MN-Is over -Ib, as expected. Thin traces are sequential single-trial sweeps from the same NMJ; the thick colored trace is the mean trace after stimulus alignment. The black curve indicates a single-exponential decay fit used to estimate τ. **(D)** Quantification of ΔR/R responses from the two inputs. **(E)** ΔF/F traces of GCaMP8m and mScarlet3 responses from single AP stimulation at MN-Ib in wild type (*w;OK319-GAL4/+;Scar8m/+*) and *GluRIIA* mutants (*w;OK319-GAL4,GluRIIA^PV3^/GluRIIA^PV3^*;*Scar8m/+*), which express presynaptic homeostatic plasticity (PHP). Note the enhanced presynaptic Ca^2+^ levels induced after PHP plasticity. **(F)** Quantification of ΔR/R responses from the indicated genotypes. Error bars represent ±SEM. Significance is indicated as *p<0.05, **p<0.01, ***p<0.001, ****p<0.0001; ns, not significant; detailed statistics including p-values are presented in Table S1.

### Evaluating the Bar8m sensor to resolve local active zone Ca^2+^ signals

Differences in Ca^2+^ influx and buffering at individual release sites contribute to functional differences at synapses (Dittman et al., 2019, Jackman et al., 2017, McCarthy et al., 2024). While Scar8m is an excellent sensor to define spatially-averaged presynaptic Ca^2+^ levels at individual boutons, the ability to resolve the heterogeneous Ca^2+^ signals within individual boutons during neurotransmission would be a powerful tool to interrogate presynaptic function. Thus, we engineered the Bar8m sensor to localize a ratiometric GCaMP8m/mScarlet3 cassette to AZs by fusion to the BRP-short motif (**Fig. 6A**). While some previous studies have suggested a similar approach can resolve AZ-specific differences in Ca^2+^ influx (Akbergenova et al., 2018), from the outset we were cognizant of many challenges to this line of investigation. First, theoretical calculations suggest that Ca^2+^ diffuses and equilibrates at a rate of <3 msec across an entire bouton, faster than most sensors would respond before receiving Ca^2+^ contributions from other release sites (Maravall et al., 2000, Sinha et al., 1997, Helmchen et al., 1997). Second, it is difficult to resolve individual AZs from live imaging of boutons, as many are closely spaced when flattened 2D images are taken. Finally, at the Drosophila NMJ, individual AZs labeled by BRP are very small, with diameters averaging ∼300 nm (Ramesh et al., 2021). To spatially resolve these discrete AZs during Ca^2+^ imaging, high spatial resolution is required, which often necessitates reducing the scanning rate. Furthermore, when employing dual-channel imaging to simultaneously capture signals from GCaMP and mScarlet, sequential scanning is typically implemented to minimize spectral crosstalk between channels. These technical constraints collectively limit the achievable frame rate for AZ-level Ca^2+^ imaging to ∼120 frames per second (fps) using resonant area scanning (Chen et al., 2024), although faster frame rates can be achieved with line scanning or CMOS cameras.

**Figure 6:**
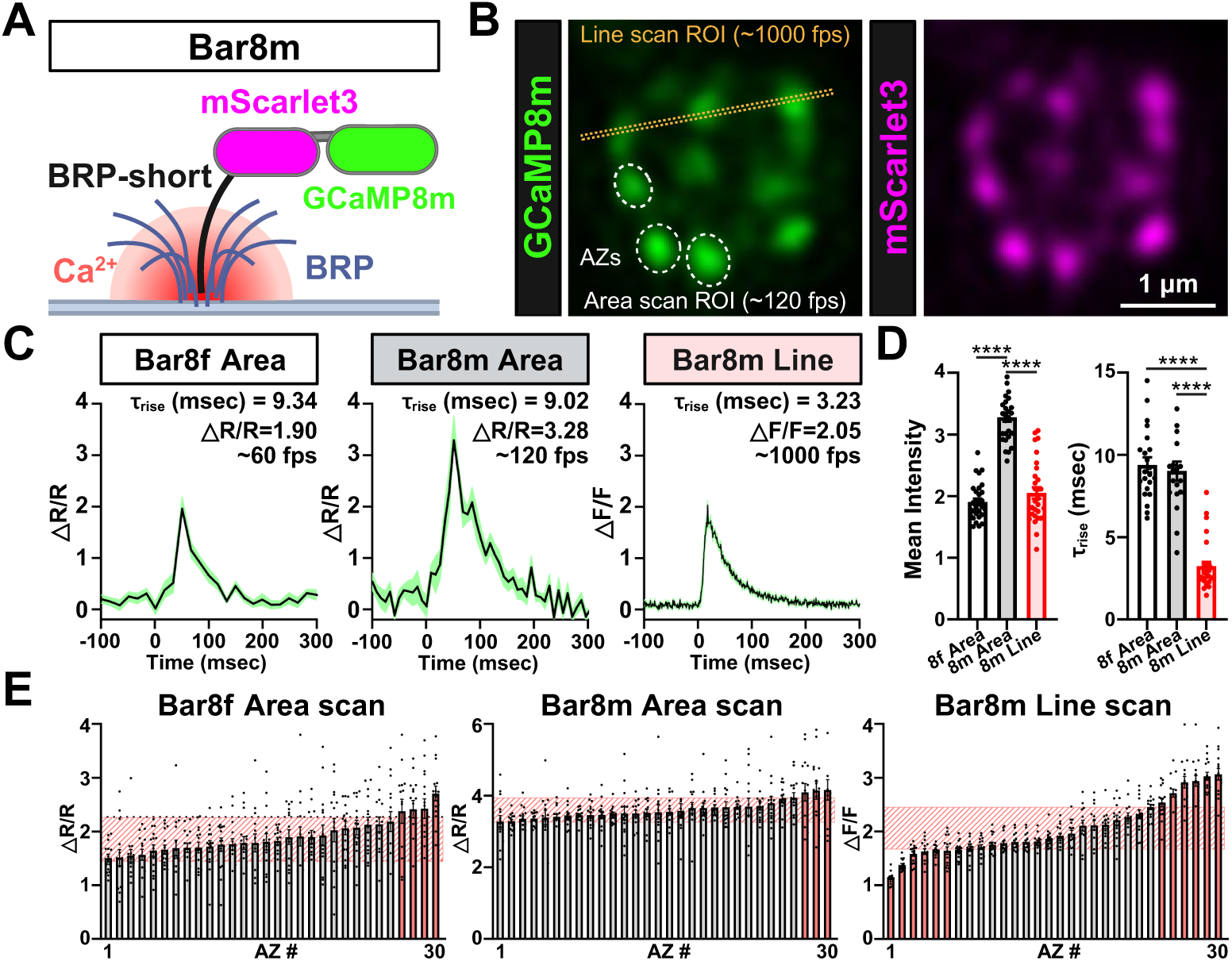
Evaluating the ability of Bar8f and Bar8m to capture active zone–specific Ca²⁺ changes. **(A)** Schematic of the BRP::mScarlet3::GCaMP8m (Bar8m) ratiometric Ca²⁺ indicator, consisting of GCaMP8m and mScarlet3 fused to the BRP-short domain, which traffics to individual AZs. **(B)** Live confocal image of a single MN-Ib bouton expressing Bar8m. Dashed circles indicate active zones used for area-scan measurements, and the yellow dotted line indicates the line-scan ROI used for high-speed imaging. **(C)** Example Ca²⁺ transients from individual AZs recorded using Bar8f resonant area scans (∼60 fps; left), Bar8m resonant area scans (∼120 fps; middle), and Bar8m line scans (∼1000 fps; right). Black traces show the mean ratiometric response (ΔR/R) for Bar8f and Bar8m area scans and the mean ΔF/F response for Bar8m line scans from a representative NMJ; green shading denotes ±SEM across repeated trials. **(D)** Quantification of mean evoked response amplitude and τ_rise_ for the three imaging conditions shown in (C). **(E)** Bar graph showing peak ΔR/R (or ΔF/F) values from individual AZs collected across multiple boutons from different NMJs for Bar8f area scans, Bar8m area scans, and Bar8m line scans. Each bar represents a single active zone (sorted by amplitude within each condition). Light red shaded boxes indicate the interquartile range (IQR; Q1–Q3) of individual data points for each active zone, and active zones whose mean responses fall outside this IQR range are highlighted in pink. Significance is indicated as *p<0.05, **p<0.01, ***p<0.001, ****p<0.0001; ns, not significant; detailed statistics including p-values are presented in Table S1.

Despite these acknowledged challenges and potential confounds, we sought to test the ability of Bar8m to resolve local AZ Ca^2+^ dynamics following single AP stimulation. Individual AZs were identified at MN-Ib boutons using basal mScarlet3 fluorescence, then evoked Ca²⁺ signals were measured using either resonant area or line scanning (**Fig. 6B**). Peak ΔR/R values were measured at individual AZs (**Fig. 6B,C**), where typical ΔR/R values at individual AZs ranged from ∼1.5 - 2.5 (Bar8f), 2.0 – 3.5 (Bar8m) using resonant area or line scanning (**Fig. 6D**). These results, while taking into account the caveats detailed above, are consistent with the theoretical estimates of Ca^2+^ levels being equilibrated across all AZs before the Ca^2+^ indicator can capture local differences, at least using relatively slow resonant area scanning approaches.

To further assess Ca^2+^ levels at individual AZ captured by the Bar8f/m sensor, we analyzed our imaging data in more detail. We plotted individual AZ ΔR/R values from lowest to highest using resonant area scanning (Bar8f and Bar8m), or line scanning (Bar8m) (**Fig. 6E**). While it was clear that the relatively slow area scanning was likely insufficient to resolve AZ-specific differences in Ca^2+^ levels, with only 3-4/30 AZs showing significant differences, fast line scanning of Bar8m was capable of capturing significant differences in 12/40 AZs in our imaging system. While we interpret the observed relative consistency of ΔR/R at individual AZs using resonant area scanning to be unable to reliably resolve local Ca^2+^ changes before equilibration within an individual bouton, faster approaches (line scanning or sCMOS cameras) might be able to detect such heterogeneity.

### SynapGCaMP8 sensors resolve quantal events at postsynaptic compartments

Finally, we evaluated the performance of SynapGCaMP6f, SynapGCaMP8f, and SynapGCaMP8m sensors to monitor postsynaptic “quantal” events at the Drosophila NMJ. These quantal events reflect the rapid ionic influxes that result from the spontaneous release of single synaptic vesicles, which open postsynaptic GluRs and allow passage of Na^+^ and Ca^2+^ ions (Grienberger et al., 2012). Electrophysiological methods are the gold standard to report these mEPSP events, and we were particularly interested in determining whether SynapGCaMP variants are capable of detecting quantal events with similar sensitivity.

We first confirmed that all SynapGCaMP variants expressed well and trafficked to postsynaptic compartments, localizing with the scaffold DLG while encompassing but being distinct from glutamate receptive fields, which is particularly apparent with super resolution STED microscopy (**Fig. 7A**). Using resonant area scanning at ∼115 fps, we imaged and analyzed single quantal Ca^2+^ events for each SynapGCaMP variant. Averaged traces revealed a progressive improvement in response kinetics and sensitivity across the indicator series (**Fig. 7B,C**). SynapGCaMP6f exhibited modest sensitivity, with ΔF/F values of 0.27 and relatively slow kinetics (τ_rise_=21 msec, τ_decay_=99 msec), while SynapGCaMP8f achieved faster responses and higher sensitivity (ΔF/F=0.35; τ_rise_=14 msec, τ_decay_=42 msec). SynapGCaMP8m exhibited the highest peak amplitude (ΔF/F=0.58), with similarly fast rise times (τ_rise_=14 msec) and moderate slowing of the decay (τ_decay_=67 msec), striking an optimal balance between speed and sensitivity (**Fig. 7C,E**). SynapGCaMP quantal signals appeared to qualitatively reflect the same events measured with electrophysiological recordings (**Fig. 7D**). We conclude that SynapGCaMP8m is an optimal indicator to measure quantal transmission events at the synapse.

**Figure 7:**
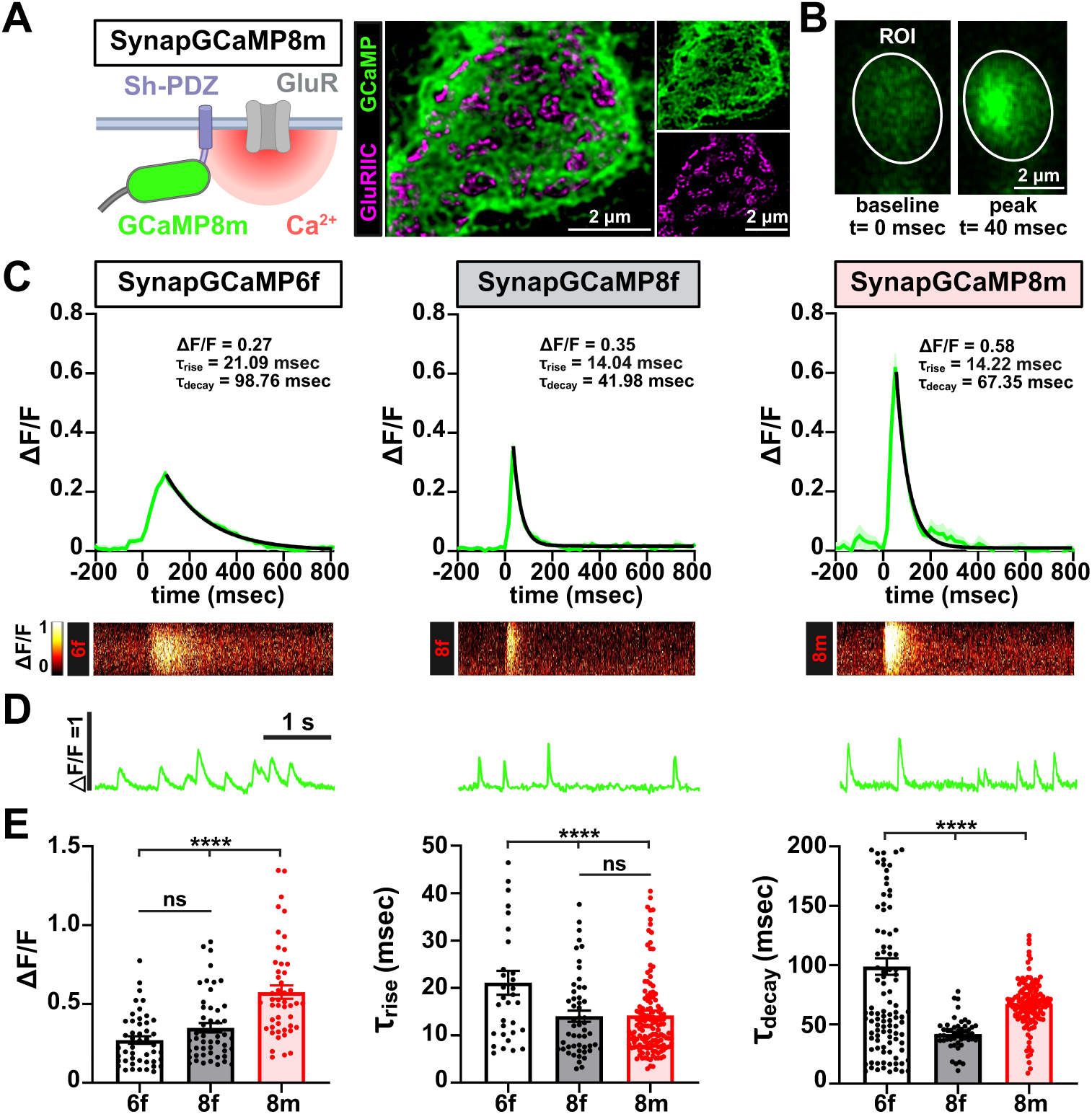
SynapGCaMP8m is an optimal postsynaptic Ca²⁺ indicator. **(A)** Schematic of the SynapGCaMP8m reporter, with GCaMP targeted to postsynaptic compartments near glutamate receptors via a Shaker-PDZ motif. Super resolution image using STED microscopy showing the GCaMP8m reporter is localized outside of GluRs. **(B)** Live confocal images of muscle 6 NMJ boutons expressing SynapGCaMP8m were performed using resonant area scans. The indicated ROI shows representative frames at baseline and peak quantal Ca²⁺ transients acquired at ∼115 fps. **(C)** Averaged single miniature Ca²⁺ events recorded from SynapGCaMP6f, -8f, and -8m. The thick green trace shows the mean ΔF/F waveform and the light green shading indicates ±SEM; the black curve over the decay phase is a single-exponential fit used to estimate τ_decay_. SynapGCaMP8m yields the highest peak signal and maintains rapid kinetics. The corresponding heatmaps below were generated from a single vertical line scan extracted from the representative miniature-event ROI, and visualize a spatiotemporal fluorescence dynamics (ΔF/F) along that line over time. **(D)** Representative Ca^2+^ events reporting individual miniature transmission in the indicated sensors. **(E)** Quantification of ΔF/F peak amplitude, and rise and decay time constants (τ_rise_ and τ_decay_) for each SynapGCaMP variant. All comparisons in bar graphs are statistically significant unless “ns” is shown. Error bars indicate ±SEM. Significance is indicated as *p<0.05, **p<0.01, ***p<0.001, ****p<0.0001; ns, not significant; detailed statistics including p-values are presented in Table S1.

Next, we systematically assessed the ability of the SynapGCaMP variants to resolve quantal events. In particular, we switched to a widefield Ca^2+^ imaging system mounted on an electrophysiology rig so that we could perform simultaneous Ca^2+^ imaging and electrophysiological recordings of mEPSP events. To simplify our analysis, we isolated miniature events from MN-Ib inputs by silencing MN-Is transmission using selective expression of BoNT-C (Han et al., 2022) (see methods). Sharp electrode intracellular recordings and Ca^2+^ imaging from muscle 6 MN-Ib boutons expressing SynapGCaMP6f, -8f, or -8m revealed increasing fidelity of quantal events (**Fig. 8A,B**). Synchronized GCaMP and mEPSP recordings showed a high degree of temporal correspondence and sensitivity between mEPSPs and Ca²⁺ transients, with SynapGCaMP8m capturing the vast majority of electrophysiologically detected mEPSP events (**Fig. 8A,B**). Indeed, mEPSP events that failed to register a detectable Ca^2+^ transient were most frequent with SynapGCaMP6f (∼40%), rare with SynapGCaMP8f (∼10%), and nearly absent with SynapGCaMP8m (∼7%) (**Fig. 8B**). Importantly, when quantal amplitudes from paired mEPSPs and corresponding Ca²⁺ transients were plotted, SynapGCaMP8m exhibited a strong linear correlation with electrophysiological recordings (Pearson’s r =0.810, R² =0.656; p<0.0001, n=216), demonstrating that the indicator reliably tracks quantal variability (**Fig. 8D**). SynapGCaMP6f and SynapGCaMP8f also showed significant positive correlations, although weaker than SynapGCaMP8m (6f: r =0.458, R²=0.210, n=283; 8f: r =0.732, R² =0.537, n=298; all p<0.0001) (**Fig. 8C**). Thus, SynapGCaMP8m provides a sensitive indicator of quantal events, approaching the sensitivity of electrophysiological recordings.

**Figure 8:**
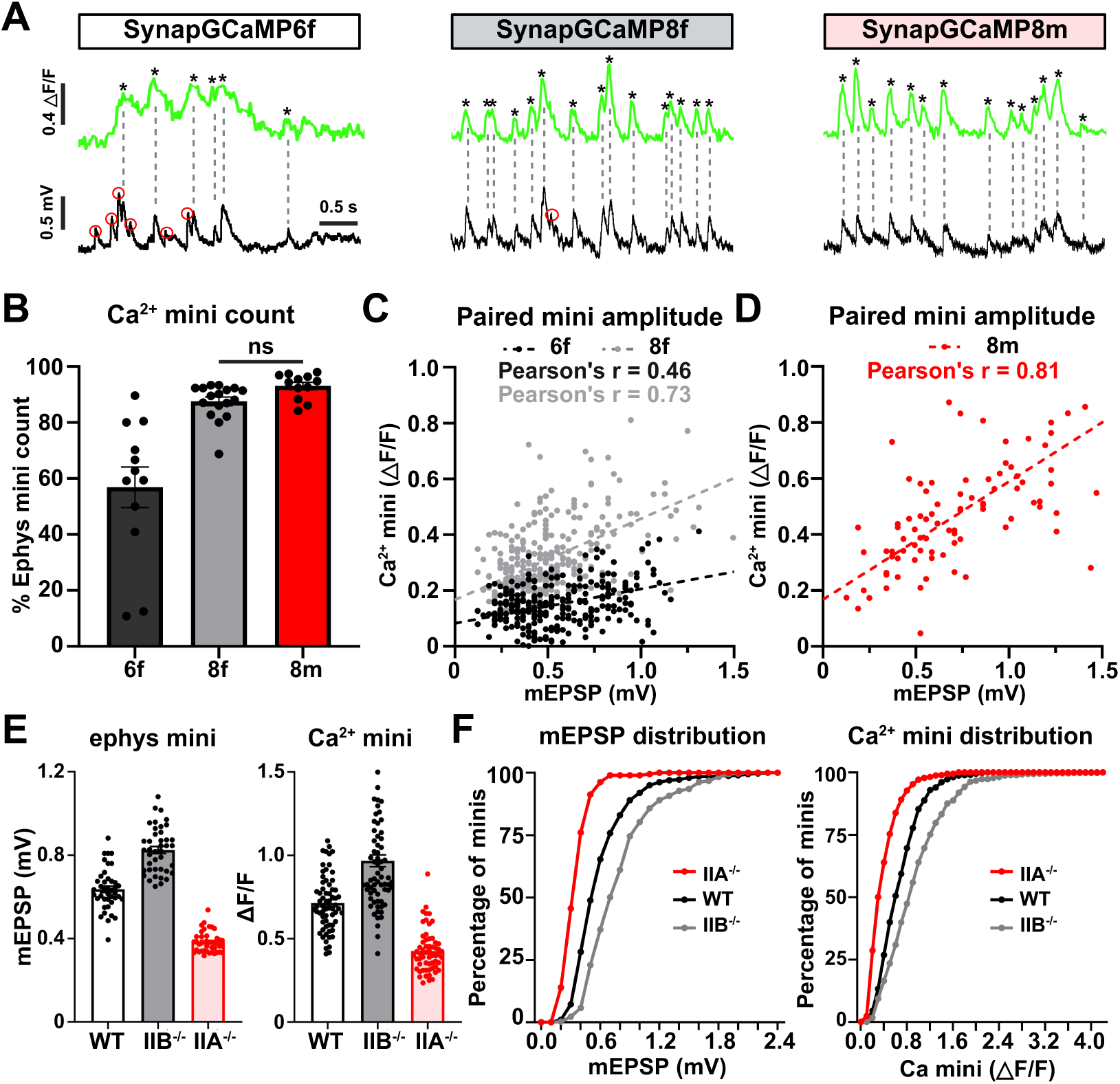
SynapGCaMP8m quantal resolution approaches that of electrophysiology. **(A)** Simultaneous recordings of quantal events at MN-Ib boutons using the indicated SynapGCaMP variant (green) and electrophysiology (black) after silencing MN-Is (*w;Is-GAL4/+;UAS-BoNT-C/SynapGCaMP*). Red circles indicate mEPSP events not captured by SynapGCaMP, and asterisks mark Ca²⁺ minis detected optically. SynapGCaMP8m captures quantal events with high sensitivity, comparable to electrophysiology. **(B)** Quantification of the proportion of quantal events captured by the indicated SynapGCaMP variant as a proportion of the total mEPSP events recorded by electrophysiology. SynapGCaMP6f detects only about half of electrophysiological events, while both SynapGCaMP8f and SynapGCaMP8m capture nearly all mEPSPs. **(C)** Scatter plot of paired miniature event amplitudes recorded simultaneously by SynapGCaMP6f and -8f and electrophysiology. Each point represents a single matched event; Pearson’s correlation coefficients (r=0.46 for 6f and r=0.73 for 8f) are indicated. **(D)** Scatter plot of paired miniature event amplitudes recorded simultaneously by SynapGCaMP8m and electrophysiology. Each point represents a single matched event; a strong linear relationship is observed (Pearson’s r=0.81), indicating that optical signals scale proportionally with quantal amplitude. **(E)** Bar plots showing average mEPSP amplitudes (left) and ΔF/F amplitudes of quantal Ca²⁺ events (right) in the indicated genotypes (same conditions as in (A) but including *GluRIIA* or *GluRIIB* mutant alleles). SynapGCaMP8m accurately resolves quantal size differences with similar resolution as the electrophysiological data, with quantal amplitudes in both datasets exhibiting the expected relationship (GluRIIA^-/-^<WT<GluRIIB^-/-^). Bars show mean ±SEM, dots represent individual boutons. **(F)** Cumulative frequency distributions of mEPSP amplitudes (left) and Ca^2+^ mini event amplitudes (right); each are significantly different using the Kolmogorov–Smirnov Test. See Table S1 for full statistical details including p-values.

In our final set of experiments, we focused on benchmarking the ability of SynapGCaMP8m to resolve physiologically relevant differences in quantal events compared to electrophysiology. Specifically, we compared electrophysiological recordings and SynapGCaMP8m imaging of quantal events in three genotypes with known differences in quantal amplitudes: wild type as the baseline control, *GluRIIA* mutants with diminished quantal amplitudes, and *GluRIIB* mutants with enlarged quantal events (Han et al., 2023).

Electrophysiological recordings from each of the three genotypes confirmed the expected differences, with baseline mEPSPs from MN-Ib averaging ∼0.64 mV, *GluRIIB* mutants ∼0.82 mV, and *GluRIIA* mutants ∼0.38 mV (**Fig. 8E**), as shown in previous studies (Han et al., 2023, Han et al., 2022, He et al., 2023). Plotting the mEPSP distribution as a cumulative probability histogram also showed the expected distribution of these genotypes (**Fig. 8F**). We then performed a similar analysis of quantal events imaged with SynapGCaMP8m. These results demonstrated near identical differences in average quantal amplitude and distribution in quantal events imaged from wild type, *GluRIIA*, and *GluRIIB* mutants (**Fig. 8E,F**). In simultaneous SynapGCaMP8m imaging and current-clamp recordings, the vast majority of optical miniature events were accompanied by a corresponding mEPSP, with less than 5% of SynapGCaMP8m transients lacking a clearly detectable mEPSP; these may reflect very small quantal events that fall below the electrophysiological detection threshold (∼0.2 mV). These data demonstrate that SynapGCaMP8m can capture physiologically relevant differences in quantal events with similar sensitivity as electrophysiology.

## DISCUSSION

This study presents a significant advancement in our capacity to optically interrogate Ca^2+^ dynamics at synaptic compartments of the Drosophila neuromuscular junction, approaches that in principle can be extended to other systems. By systematically engineering and rigorously benchmarking a new suite of ratiometric GECIs – presynaptic Scar8m, active-zone targeted Bar8m, and postsynaptic SynapGCaMP8m – and complemented with the development of the CaFire automated analysis platform, we have overcome limitations of previous tools. These indicators provide a powerful toolkit for dissecting the Ca^2+^ signals that mediate and control synaptic transmission and plasticity, moving us closer to all optical interrogation with similar resolution as electrophysiology. These results demonstrate that careful sensor design, incorporating the latest GCaMP8m variant and optimized ratiometric partners like mScarlet3, can yield probes with unprecedented performance in this model system.

Presynaptic Ca^2+^ imaging with Scar8m not only surpasses previous GECIs but also rivals, and in some respects exceeds, the capabilities of traditional synthetic Ca^2+^ indicators. While chemical dyes offer rapid kinetics (Hendel et al., 2008, Lock et al., 2015, Macleod, 2012, Lnenicka et al., 2006, Justs et al., 2022, Lu et al., 2016), their utility is often hampered by inconsistent loading/concentrations, lack of cell-type specificity, and challenges in ratiometric quantification within the confined space of presynaptic boutons. Indeed, traditional approaches to load dyes into fly motor neurons relies on cutting the nerve while bathed in solution, and waiting for the dye to diffuse the considerable distance to presynaptic terminals, where variation in dye concentration can confound analyses (McMahon et al., 2018, Macleod, 2012). Scar8m, targeted to synaptic vesicle pools at boutons via fusion to Synaptotagmin along with the ratiometric mScarlet3 motif, circumvents these issues, providing robust, genetically targeted, and stoichiometric and quantifiable Ca^2+^ measurements, spatially-averaged and confined to individual boutons that outperform cytosolic GCaMPs (Fig. 3). Further, the superior sensitivity and relatively fast kinetics of Scar8f/m enabled the resolution of physiologically meaningful differences in presynaptic Ca^2+^ levels, such as those between MN-Ib and MN-Is terminals and the elevations associated with presynaptic homeostatic plasticity (Li et al., 2021). When directly comparing OGB-1 to Scar8f, we found superior sensitivity and kinetics (Fig. 3), which, by extrapolation, should also compare favorably to other synthetic dyes like rhod and Fura (Lu et al., 2016, He et al., 2023). Notably, Scar8f achieves a ΔF/F of approximately 0.32 vs 0.23 for OGB-1, while also showing a faster decay time constant (∼40 msec for Scar8f vs. ∼60 msec for OGB-1). Furthermore, the rise time performance of Scar8f/m is on par with that of OGB-1, with a single-AP rise time of ∼12 msec compared to ∼8 msec (Fig. 3), with the caveat that scan speed can limit the accuracy of rise times. This capacity to faithfully report presynaptic Ca^2+^ changes, previously accessible primarily via chemical dyes, underscores Scar8m’s power as a tool for investigating presynaptic function with enhanced precision and reliability.

Equally significant is the performance of the postsynaptically targeted SynapGCaMP8m, which we have shown achieves a sensitivity for detecting quantal Ca^2+^ events that approaches the gold standard of electrophysiological recordings. Previous optical methods often struggled to reliably capture these small, stochastic events, particularly in the case of glutamate receptor loss or perturbation (Newman et al., 2017, Peled et al., 2014, Han et al., 2022), and have never previously been benchmarked simultaneously with electrophysiological recordings to determine their fidelity. We found that SynapGCaMP6f failed to capture over 40% of mEPSP events, while the GCaMP8m version reliably detected over 90% of these events, even in cases of diminished amplitude due to *GluRIIA* loss. Leveraging the high responsivity of GCaMP8m enabled substantially improved peak amplitudes and kinetics. Furthermore, the strong correlation observed between the amplitudes of optical quantal events and their corresponding mEPSPs, along with the ability to accurately resolve known differences in quantal size in *GluRIIA* and *GluRIIB* mutants, serves to validate SynapGCaMP8m as a high-fidelity reporter of postsynaptic activity. Finally, we note the additional benefits of quantal imaging when compared to electrophysiology, where spatial discrimination of quantal events allows, for example, the ability to quantify input-specific events (see **Supplemental Video 1**). This opens exciting avenues for all-optical interrogation of quantal parameters, synaptic strength, and changes due to plasticity with subcellular spatial resolution.

Our investigation into capturing local Ca^2+^ signals using an active zone-targeted, ratiometric GCaMP8m based indicator, Bar8m, while ambitious, also yielded important insights despite the inherent technical challenges. Bar8m was able to sensitively report Ca^2+^ changes at individual AZs following single AP electrical stimulation, as evidenced by the significant differences observed across AZs, expected by variation in Ca_v_2 channel abundance shown in previous studies(Cunningham et al., 2022, Gratz et al., 2019, Medeiros et al., 2024). However, the Bar8f/m intensity signals showed relative consistency across AZs following single AP stimulation and imaged using resonant area scans. This finding, coupled with theoretical considerations of Ca^2+^ diffusion rates (Sinha et al., 1997, Helmchen et al., 1997), suggests that within the temporal resolution of current GCaMP sensors, and likely synthetic dyes, Ca^2+^ concentrations are likely to equilibrate across AZs within an individual bouton before distinct local concentration differences can be reliably resolved. There are a number of other potential confounds precluding a definitive conclusion about the ability of Bar8f/m to resolve local Ca^2+^ changes, including challenges in resolving individual AZs and imaging speeds. Indeed, more promising results were observed using line scanning, with kHz speeds, of Bar8m, which appeared to capture a higher degree of AZ-specific differences (Fig. 6E). These observations are critical for interpreting data acquired from AZ-targeted GECIs and guide future efforts to capture localized Ca^2+^ nanodomain dynamics, likely requiring sensors with even faster kinetics and/or alternative imaging modalities.

The suite of next-generation GECIs, particularly Scar8m and SynapGCaMP8m, coupled with the CaFire analysis pipeline, represents a substantial toolkit for studying synaptic Ca²⁺ signaling. At the Drosophila NMJ, these tools provide researchers with unprecedented optical access to both presynaptic Ca^2+^ influx driving neurotransmitter release and postsynaptic Ca²⁺ transients reflecting quantal events, with performance characteristics that rival traditional, more invasive methods. The ability to perform ratiometric, genetically-targeted imaging with such high sensitivity and temporal resolution will help to accelerate discoveries into the molecular mechanisms of synaptic transmission, plasticity, and disease etiology. More generally, the strategies used here in the fly system can inspire similar approaches to be employed in other systems. Future studies leveraging these advanced sensors can now tackle complex questions regarding the spatial and temporal dynamics of Ca^2+^ signaling within synaptic compartments, the modulation of these signals during various forms of plasticity, and their dysregulation in models of neurological disorders, further cementing the Drosophila NMJ as a premier system for fundamental synaptic research.

## MATERIALS AND METHODS

### Fly stocks

Drosophila stocks were raised at 25°C using standard molasses food. Unless otherwise specified, the *w^1118^* strain was used as the wild-type control as this is the genetic background in which all genotypes were bred. All experiments were performed on Drosophila third-instar larvae of both sexes unless otherwise noted. See Table S2 (Key Resources Table) for a full list of all fly stocks and their sources used in this study.

### Molecular biology

To generate the transgenic constructs used in this study, we built upon previously established tools. First, SynapGCaMP8m was engineered by modifying the previous SynapGCaMP8f construct (Han et al., 2022), which was in turn based on the original SynapGCaMP6f construct (Newman et al., 2017). Gibson assembly reactions were used to convert the few relevant amino acid differences between GCaMP8f and GCaMP8m (Zhang et al., 2023). Similarly, UAS-Syt::mScarlet3::GCaMP8m (Scar8m) was generated using similar approaches to modify the previously described UAS-Syt::mScarlet::GCaMP8f plasmid (Li et al., 2021), changing both the GCaMP8f and mScarlet sequences to GCaMP8m and mScarlet3 (Gadella et al., 2023). Finally, UAS-BrpS::mScarlet::GCaMP8f (Bar8f) and UAS-BrpS::mScarlet3::GCaMP8m (Bar8m) were engineered based on the previous UAS-BrpS::mCherry::GCaMP6s transgene (Kiragasi et al., 2017), where the GCaMP6s sequence was replaced with GCaMP8f or GCaMP8m, and the mCherry tag was substituted with mScarlet or mScarlet3. SynapGCaMP transgenes were inserted randomly through p-element transposition, while all other transgenes were inserted into the attP40 (II) or VK27 (III) sites by BestGene Inc. (Chino Hills, CA). All constructs were verified by Sanger sequencing prior to injection, and transgenic fly stocks were established and maintained under standard laboratory conditions.

### Immunocytochemistry

Third-instar larvae were dissected in ice cold 0 Ca^2+^ HL-3 and immunostained as described (Li et al., 2021, Han et al., 2023). Briefly, larvae were either fixed in 100% ice-cold methanol for 5 mins or PFA for 10 mins followed by washing with PBS containing 0.1% Triton X-100 (PBST). Samples were blocked with 5% Normal Donkey Serum for 1 hour and incubated with primary antibodies overnight at 4°C. Preparations were washed for 10 mins thrice in PBST, incubated with secondary antibodies for 2 hours at room temperature, washed thrice again in PBST, and equilibrated in 70% glycerol. Prior to imaging, samples were transferred in VectaShield (Vector Laboratories, Burlingame, CA) and mounted on glass cover slides. Details of all antibodies and sources are listed in Table S2.

### Electrophysiology

All dissections and electrophysiological recordings were performed as described (Kikuma et al., 2019) in modified hemolymph-like saline (HL-3) containing (in mM): 70 NaCl, 5 KCl, 10 MgCl_2_, 10 NaHCO_3_, 115 Sucrose, 5 Trehelose, 5 HEPES, pH=7.2, and CaCl_2_ at 0.4 mM unless otherwise specified. Internal guts, brain and the ventral nerve cord were subsequently removed to achieve fully dissected preparations. Recordings were carried out on an Olympus BX61 WI microscope stage equipped with a 40x/0.8 NA water-dipping objective and acquired using an Axoclamp 900A amplifier (Molecular Devices). All recordings were conducted on abdominal muscle 6, segment A3 of third-instar larvae of both sexes. Data were acquired from cells with an initial resting potential between -60 and -80 mV, and input resistances >5 MΩ. Miniature excitatory postsynaptic potentials (mEPSPs) were recorded without any stimulation and low pass filtered at 1 kHz. The mEPSPs for each sample were recorded for 60 secs, analyzed with MiniAnalysis (Synaptosoft), and the average mEPSP amplitude for each NMJ was calculated. Excitatory postsynaptic potentials (EPSPs) were recorded by delivering 20 electrical stimuli at 0.5 Hz with 0.5 msec duration to motor neurons using an ISO-Flex stimulus isolator (A.M.P.I.) with stimulus intensities set to avoid eliciting multiple EPSPs.

### Confocal imaging

Dissections and live Ca^2+^ imaging was performed as described (Chien et al., 2025, Chen et al., 2024) on muscle 6 NMJs. Wandering third-instar larvae were dissected and imaged in 1.8 mM Ca^2+^ HL3 saline. Ca^2+^ imaging was conducted using a Nikon A1R resonant scanning confocal microscope equipped with a 60x/1.0 NA water-immersion objective (refractive index 1.33). GCaMP signals were acquired using the FITC/GFP channel (488-nm laser excitation; emission collected with a 525/50-nm band-pass filter), and mScarlet/mCherry signals were acquired using the TRITC/mCherry channel (561-nm laser excitation; emission collected with a 595/50-nm band-pass filter). ROIs focused on terminal boutons of MN-Ib or -Is motor neurons. For both channels, the confocal pinhole was set to a fixed diameter of 117.5 µm (approximately three Airy units under these conditions), which increases signal collection while maintaining adequate optical sectioning. Images were acquired as 256 × 64 pixel frames (two 12-bit channels) using bidirectional resonant scanning at a frame rate of ∼118 frames/s; the scan zoom in NIS-Elements was adjusted so that this field of view encompassed the entire neuromuscular junction and was kept constant across experiments. In ratiometric recordings, the 488-nm (GCaMP) and 561-nm (mScarlet) channels were acquired in a sequential dual-channel mode using the same bidirectional resonant scan settings: for each time point, a frame was first collected in the green channel and then immediately in the red channel, introducing a small, fixed frame-to-frame temporal offset while preserving matched spatial sampling of the two channels. Resonant area scans of Bar8f and Bar8m responses were acquired at ∼60 or ∼120 frames/s with scanning area of 256 × 32–64 pixels, using a zoom factor of 8×. Because Bar8f was dimmer, the confocal image quality setting was increased from level 1 to 2, which reduced the scanning frame rate. To resolve rapid Ca²⁺ transients at individual AZs, we additionally performed single-channel Galvano line-scan imaging of Bar8m. A brief resonant area image was first acquired to localize BRP-positive puncta, and a one-dimensional scan line was then positioned through the center of each selected AZ. Line-scan images were acquired at ∼1000 frames/s sampling rate. Presynaptic and active zone Ca^2+^ imaging was conducted for 15 secs per acquisition while delivering electrical stimulation at 1 Hz (1 msec pulse duration) or at 5 Hz and 10 Hz (five pulses per 2-second burst). Action potentials were evoked by extracellular stimulation of the segmental motor nerve using a glass suction electrode. Suction electrodes were pulled from TW120-4 glass capillaries on a Sutter P-97 puller and fire-polished to a final tip opening of ∼5-10 µm, then filled with HL3 saline. The cut motor nerve innervating the imaged muscle was gently drawn into the electrode, and electric stimulation was delivered through a pulse stimulator coupled to an isolated stimulator (Iso-Flex, controlled by a Master-9 pulse generator)(Chen et al., 2024). To prevent muscle contraction during imaging, 7 mM L-Glutamic acid monosodium salt (Sigma-Aldrich, Cat# G5889) was added to the HL3 saline. In contrast, postsynaptic Ca^2+^ imaging was performed for 60 secs without electrical stimulation, as only quantal events were monitored. Data was collected from at least three biological replicates per genotype.

Time-lapse videos were stabilized and bleach-corrected prior to analysis, which visibly reduced frame-to-frame motion and intensity drift. In the presynaptic and active-zone mScarlet channel, a bleaching factor of ∼1.15 was observed during the 15 sec recording. This bleaching can be corrected using the “Bleaching correction” tool in Huygens SVI. For presynaptic and active-zone GCaMP signals, there was minimal bleaching over these short imaging periods.

Therefore, the bleaching correction step for GCaMP was skipped. Both GCaMP and mScarlet channels were processed using the default settings in the Huygens SVI “Deconvolution Wizard” (with the exception of the bleaching correction option). Deconvolution was performed using the CMLE algorithm with the Huygens default stopping criterion and a maximum of 30 iterations, such that the algorithm either converged earlier or, if convergence was not reached, was terminated at this 30-iteration limit; no other iteration settings were used across the GCaMP series. ROIs were drawn on the processed images using Fiji ImageJ software, and mean fluorescence time courses were extracted for the GCaMP and mScarlet channels, yielding F_GCaMP_(t) and F_mScarlet_(t). F(t)s were imported into CaFire with GCaMP assigned to Channel #1 (signal; required) and mScarlet to Channel #2 (baseline/reference; optional). If desired, the mScarlet signal could be smoothed in CaFire using a user-specified moving-average window to reduce high-frequency noise. In CaFire’s ΔR/R mode, the per-frame ratio was computed as R(t)=F_GCaMP_(t)/F_mScarlet_(t); a baseline ratio R0 was estimated from the pre-stimulus period, and the final response was reported as ΔR/R(t)=[R(t)−R0]/R0, which normalizes GCaMP signals to the co-expressed mScarlet reference and thereby reduces variability arising from differences in sensor expression level or illumination across AZs (see CaFire section in methods for details).

Confocal images of fixed tissue were acquired with a 100x APO 1.45NA oil immersion objective using separate channels with four laser lines (405 nm, 488 nm, 561 nm, and 647 nm) as described (Perry et al., 2017). Z-stacks were obtained on the same day using identical gain and laser power settings with z-axis spacing between 0.13 and 0.2 µm and pixel size of 0.06 µm for all samples within an individual experiment, from at least 8 NMJs acquired from at least four different animals. Raw confocal images were deconvolved with SVI Huygens Essential 22.10 using built-in Express settings. All confocal images shown in the figures, including Fig. 1, were deconvolved in Huygens using this CMLE-based workflow prior to maximum-intensity projection and display. Maximum intensity projections were created for quantitative image analysis using the general analysis toolkit of NIS Elements software.

### Benchmarking OGB-1 vs Scar8f

For OGB-1 imaging, wild-type motor neuron terminals were forward filled with Oregon Green 488 BAPTA-1 (OGB-1; 10,000 MW anionic dextran, batch-specific Kd = 1180 nM, Molecular Probes) as described (Macleod et al., 2002). Larvae were rinsed in distilled water, transferred to Schneider’s Drosophila medium, and pinned dorsal-side up on Sylgard-coated dishes with Minutien pins. A midline incision was made along the dorsal surface, the body wall was pinned open to generate a fillet preparation, and internal organs were removed to expose the ventral nerve cord and segmental nerves. For forward filling, segmental nerves were severed near the ventral ganglion and the cut ends from hemisegment 4 were drawn into a glass suction pipette; a bolus of 5 mM OGB-1 in distilled water was ejected onto the nerve ends in an equal volume of Schneider’s medium, left in place for 30–60 min, then replaced with Schneider’s medium in the pipette to serve as the cathode for stimulation. OGB-1 was allowed to transport to NM6/7-Ib terminals for 4–5 h at room temperature, with Schneider’s medium exchanged hourly. Immediately prior to imaging, Schneider’s medium was replaced with hemolymph-like 6 (HL6) saline containing 2 mM CaCl₂ and 7 mM L-glutamic acid for at least 15 min. Ca^2+^ imaging was performed on an upright confocal laser-scanning microscope (Olympus Fluoview 4000) equipped with a LUMPlanFl 60×/0.90 W objective. GCaMP8f or OGB-1 were excited at 488 nm using 10% and 5% laser power, respectively, and emission was collected through a maximized pinhole using a grating and slit set to 510–550 nm. mScarlet was excited at 561 nm (2% power) and emission collected at 570–620 nm. Images were acquired with a 2× digital zoom (190 nm/pixel) over a 512×48 pixel frame at 3.295 msec per frame. For each recording, 1 sec of baseline was collected prior to nerve stimulation, followed by 14 sec of imaging to capture 10 isolated single AP responses and a subsequent train of 17 APs. Nerves innervating segment 4 were stimulated using the silver wire in the suction pipette as the cathode and a bath wire as the anode, driven by an isolated pulse stimulator (DS2A, Digitimer) controlled by a Master-8 (A.M.P.I.). Stimuli were 1.5 V, 0.3 msec pulses delivered as 10 APs at 1 Hz, followed after a 2 sec delay by 17 APs at 21.3 Hz. Only preparations with stable baseline fluorescence and reproducible responses were analyzed. For each indicator, 7 larvae were imaged, with the full stimulus trial (10 @ 1 Hz followed by 17 @ 21.3 Hz) repeated three times per larva. Fluorescence was quantified from ROIs surrounding 1–2 non-terminating MN6/7-Ib boutons; background-subtracted stimulus fluorescence (F_stim_) was obtained using cellSens FV software (v4.1.1, Evident Scientific) and further processed in Microsoft Excel. Bleaching (typically ∼8 % and 12 % per 10 sec for GCaMP8f and OGB-1, respectively) was corrected by fitting a linear trend to the average pre-stimulus fluorescence in each 100 msec window preceding the 10 APs at 1 Hz and adjusting each F_stim according to its time and the slope of this line. ΔF/F was then calculated as (F_stim_-F_rest_)/F_rest_, where F_rest_ is the mean fluorescence during the 100 msec preceding the first AP. mScarlet fluorescence was not used when benchmarking Scar8f against OGB-1 because a red dextran fluorophore was not co-loaded in constant proportion with OGB-1. For single-AP analysis, three trials per preparation were averaged, and the 10 responses were aligned so that the first rise in F_stim_ occurred 1.648 msec (half a sampling interval) after the stimulus. Amplitude was obtained from the intercept at time 0 of a single-exponential fit (ΔF/F = ΔF/F=A_o_e^-bt^) to data from 14.828–499.193 msec (147 points), and the decay time constant τ was calculated as 1000 ms/−b. For train responses (17 @ 21.3 Hz), trials were averaged after alignment to the first rise in F_stim_; amplitude was taken as the peak ΔF/F after the final AP, the facilitation index was computed as the ratio of the final to the first response, and τ was derived from a single-exponential fit to 1 sec (304 points) of data following the peak.

### Widefield Ca^2+^ imaging

Widefield Ca^2+^ imaging was conducted on muscle 4 NMJs using a Zeiss Axio Examiner A1 upright fixed-stage microscope equipped with a pco.panda 4.2 sCMOS camera (Excelitas) in 1.8 mM Ca^2+^ HL3 saline. High-power 470 nm LED (Thorlabs, M470L4) was used for illumination with a 63×/1.0 NA water-dipping lens, with image acquisition controlled using Nikon NIS-Elements software. Regions of interest (ROIs) were defined to encompass terminal boutons of MN-Ib branches on muscle 4. Images were acquired at 256×256 pixels with a frame rate of 100 fps. GCaMP was imaged using a GFP filter cube (LED excitation ∼470/40 nm, emission ∼525/50 nm). Time-series imaging was performed for 60 secs without external stimulation to capture spontaneous quantal Ca²⁺ events. Under these widefield imaging conditions, SynapGCaMP signals exhibited only modest photobleaching, typically a ∼2–5% decrease in baseline fluorescence over a 60 sec acquisition. The modest bleaching was either corrected using the “bleaching correction” function in Huygens SVI or accounted for during ΔF/F quantification in the CaFire analysis pipeline using a sliding-baseline procedure (see CaFire methods). For simultaneous electrophysiological recordings, Clampex software (Molecular Devices) was employed to perform current-clamp recordings in gap-free mode. A digital TTL trigger signal from Clampex synchronized the imaging acquisition in NIS-Elements, ensuring precise temporal alignment between electrophysiological and imaging data. Data was collected from a minimum of six biological replicates per genotype.

### STED imaging

STED super-resolution microscopy was performed as previously described(He et al., 2023, Chien et al., 2025). Briefly, STED imaging was performed with an Abberior STEDYCON system mounted on a Nikon Eclipse FN1 upright microscope. The system is equipped with four excitation lasers (640, 561, 488 and 405 nm), a pulsed STED laser at 775 nm, and three avalanche photodiode detectors operating in a single photon counting mode.

Alexa Fluor 488-conjugated anti-HRP was used to locate NMJs, whereas Abberior STAR Red and Alexa Fluor 594 secondary dyes were used for the STED channels. Depletion was performed at 775 nm with time-gated detection set to open from 1 ns to 7 ns after each excitation pulse. Emission was detected with two avalanche photodiodes (band-pass filters: 600 ± 25 nm for STAR RED and 675 ± 25 nm for Alexa Fluor 594). Images were acquired sequentially with a pixel dwell time of 10 µs and 3× line accumulation. Multichannel z-stack STED images were acquired using a 100× Nikon Plan APO 1.45 NA oil immersion objective with a 20-nm fixed pixel size at 130 nm steps, which yields an effective lateral resolution of ∼60 nm. Each image covered one to two boutons (25-64 μm^2^) on muscle 4 of segment A3. Raw STED images were corrected for thermal drift and channel crosstalk, then deconvolved in SVI Huygens software Essential 24.04.0 (Scientific Volume Imaging B.V.) using theoretical STED point-spread and default iteration settings of the Good’s MLE algorithm.

### CaFire program

CaFire is a software tool designed for the analysis and processing of Ca^2+^ imaging data, compatible with both raw intensity data and pre-normalized signals (e.g., ΔF/F or ΔR/R). The graphical user interface (GUI) of CaFire is implemented with the Python Tkinter library, while the visualization panel is powered by Matplotlib. The method of automated peak detection utilizes signal processing functions from SciPy (Virtanen et al., 2020). We define SNR operationally as the fractional fluorescence change (ΔF/F). Peaks that are not detected automatically can be selected manually. CaFire identifies the nearest peak to the location where the user clicks. For peak quantification analysis, CaFire calculates rise and decay properties by performing curve fitting on data preceding and following the peak, respectively. The curve fitting is performed using the Levenberg–Marquardt gradient descent algorithm, implemented in the curve fit function of SciPy (Virtanen et al., 2020). The rise time constant is calculated by applying an exponential fit to the rise phase (Shemesh et al., 2020). The decay is modeled using a natural exponential decay function: *y* == *y_peak_· e* (Fleming et al., 2021).

After motion correction/processing, ROIs were drawn in Fiji and mean-intensity time series were exported as numeric tables. The traces were organized in an Excel (.xlsx) file, with the first row containing column headers (text labels) and all subsequent rows containing numeric data only. CaFire directly imports .xlsx tables and requires at least two columns per trace: time (or frame number) on the x-axis and intensity on the y-axis (additional ROIs can be placed in separate columns with distinct headers). If only frame numbers are provided, rise and decay tau results need to be converted to time separately by multiplying the fitted tau (in frames) by the frame interval (Δt=1000 msec/frame rate). CaFire supports both single- and dual-channel imaging: for all analyses, GCaMP traces are imported as Channel #1 (signal; required), and for dual-channel datasets the mScarlet reference is additionally imported as Channel #2 (baseline/reference; optional). For single-channel recordings, CaFire can either accept user-supplied ΔF/F traces or compute ΔF/F from raw fluorescence: in the latter case, CaFire first estimates a time-varying baseline F₀(t) using a sliding-window percentile method, with the window length, percentile, and time range user-configurable in the GUI (unless otherwise noted, we used a 101-point window, i.e., 50 frames before and after each time point, and the 30th percentile of values within that window to define F₀). Fluorescence signals are then expressed as ΔF/F(t) = [F(t) − F₀(t)] / F₀(t), which we report throughout the manuscript as an operational measure of signal-to-noise ratio (SNR). For ratiometric datasets, CaFire allows users to analyze either precomputed ΔR/R traces or to generate ΔR/R from raw two-channel fluorescence data. ROIs are drawn in Fiji/ImageJ and mean fluorescence time courses are extracted for the green GCaMP channel and the red mScarlet channel, yielding F_GCaMP_(t) and F_mScarlet_(t), which are imported into CaFire as the signal and reference channels, respectively. If desired, the mScarlet signal can be smoothed in CaFire using a user-specified moving-average window (e.g., 8 frames at ∼118 fps) to reduce high-frequency noise without altering the underlying ratio dynamics. In ΔR/R mode, CaFire computes a per-frame ratio R(t) = F_GCaMP_(t) / F_mScarlet_(t) and estimates a baseline ratio R₀(t) using the same sliding-window percentile procedure; for Bar8m active-zone analysis, we used a 101-point window and the lowest 10% of ratio values within each window to define R₀. Ratiometric responses are then reported as ΔR/R(t) = [R(t) − R₀(t)] / R₀(t). CaFire can plot raw, ΔF/F, or ratiometric traces, and F₀(t) or R₀(t) can be visualized in the raw-data view for quality control. The source code and a Windows executable, together with additional instructions (including ImageJ workflow instructions, an Excel template, CaFire loading screenshots, and a brief step-by-step usage guide), are available for download on CaFire’s GitHub repository (https://github.com/linj7/CaFire) and has been archived with a DOI at Zenodo: 10.5281/zenodo.1552996. The software is released under the MIT License.

### Statistical analysis

Data were analyzed using NIS Elements software (Nikon), MiniAnalysis (Synaptosoft), SVI Hugyens Essential (version 22.10), CaFire (https://github.com/linj7/CaFire), GraphPad Prism (version 10.0), and Microsoft Excel software (version 16.22). For comparisons between two groups, we used unpaired two-tailed Student’s t-tests. For comparisons among more than two groups, we used one-way ANOVA followed by Tukey’s multiple comparisons test. Cumulative distributions of mEPSP amplitudes and Ca²⁺ mini amplitudes were compared using two-sample Kolmogorov–Smirnov tests. Pearson correlation coefficients and linear regression were used to assess relationships between optical and electrophysiological miniature event amplitudes, as well as between AZ size and Ca²⁺ signal amplitude. When outlier analysis was required (e.g., analyzing AZs), outliers were identified using the interquartile range (IQR) method. To quantify AZ-to-AZ variability, we performed paired two-tailed t-tests between every pair of active zones within the same preparation, comparing peak responses across stimuli. For Bar8m line-scan recordings, ∼79% of AZ pairs were significantly different at α < 0.05, whereas for Bar8f area-scan recordings only ∼8% of AZ pairs were significant, indicating substantially greater AZ-to-AZ diversity when measured at high temporal resolution. For visual marking of outlying AZs in the scatter plots, we computed the interquartile range (IQR; Q1–Q3) from the individual responses pooled across AZs (15 stimuli per AZ, 30 AZs per genotype) and highlighted AZs whose mean response lay above Q3 or below Q1. The IQR was used solely as a robust measure of dispersion for visualization, and not as a formal hypothesis test. The fraction of significant AZ pairs is reported on the corresponding plots. Throughout the manuscript, n refers to the number of animals (biological replicates). All Ca²⁺ imaging and electrophysiological recordings were performed at abdominal muscle 6, segment A3 of third-instar larvae. For imaging experiments, each larva contributed 1–2 NMJs in total; at each NMJ we typically imaged 2–3 terminal boutons and acquired 2–3 time-series recordings from different boutons. For the standard stimulation protocol, 1-msec pulses were delivered at 1 Hz during a 15-sec imaging epoch (yielding ∼14–15 stimuli per trial). In all figures, error bars indicate ±SEM, with the following statistical significance: p<0.05 (*), p<0.01 (**), p<0.001 (***), p<0.0001 (****); ns=not significant. Additional statistical details for all experiments are summarized in Table S1.

## AUTHOR CONTRIBUTIONS

J.C. and D.D. designed the research; J.L. developed the CaFire software; J.C., J.L., K.H., L.W., Y.H., C.Q., J.W., C.D., and G.M. performed experiments and analyzed the data. The manuscript was written by J.C. and D.D. with feedback from the other authors.

## Data and Material Availability

All acquired data are available upon request to the lead contact. All fly stocks and molecular constructs generated in this study will be shared upon request. We are also in the process of depositing newly generated Scar8f/m, Bar8f/m, and SynapGCaMP sensors to the Bloomington Drosophila Stock Center for public dissemination.

The source code for the CaFire analysis software is publicly available on GitHub at https://github.com/linj7/CaFire and has been archived with a DOI at Zenodo: 10.5281/zenodo.1552996.

## ACKNOWLEDGEMENTS

We thank Greg Macleod (Tulane University, New Orleans, LA, USA), Martin Muller (University of Zurich, Zurich, Switzerland), and Igor Delvendahl (University of Freiburg, Freiburg, Germany) for useful discussions about Ca^2+^ imaging and interpretations. We acknowledge the Developmental Studies Hybridoma Bank (Iowa, USA) for antibodies used in this study and the Bloomington Drosophila Stock Center for fly stocks (NIH P40OD018537). This work was supported by grants from the National Institutes of Health NINDS awards to G.M. (NS1233777) and D.D. (NS091546 and NS126654). BioRender was used to create schematics shown in Figures 1 and 3–8, as well as Figure S1.

## Declaration of generative AI and AI-assisted technologies in the writing process

During the preparation of this work the authors used Google Gemini Advanced 2.5 Pro to line edit an early version of the manuscript to improve clarity and readability. After using this tool, the authors subsequently reviewed and edited the content and take full responsibility for the content of the publication.

## REFERENCES

Berridge MJ, Bootman MD, Roderick HL. (2003) Calcium signalling: dynamics, homeostasis and remodelling. Nature Reviews Molecular Cell Biology.47:517–29

Clapham DE. (2007) Calcium signaling. Cell.1316:1047–58

Dittman JS, Ryan TA. (2019) The control of release probability at nerve terminals. Nat Rev Neurosci.203:177–86

Jackman SL, Regehr WG. (2017) The Mechanisms and Functions of Synaptic Facilitation. Neuron.943:447–64

McCarthy CI, Kavalali ET. (2024) Nano-organization of synaptic calcium signaling. Biochemical Society Transactions.523:1459–71

Nakai J, Ohkura M, Imoto K. (2001) A high signal-to-noise Ca²⁺ probe composed of a single green fluorescent protein. Nature Biotechnology.192:137–41

Tian L, Hires SA, Mao T, Huber D, Chiappe ME, Chalasani SH, Petreanu L, Akerboom J, McKinney SA, Schreiter ER, et al. (2009) Imaging neural activity in worms, flies and mice with improved GCaMP calcium indicators. Nature Methods.612:875–81

Dana H, Mohar B, Sun Y, et al. (2019) High-performance calcium sensors for imaging activity in neuronal populations and microcompartments. Nature Methods.167:649–57

Chen TW, Wardill TJ, Sun Y, et al. (2013) Ultrasensitive fluorescent proteins for imaging neuronal activity. Nature.4997458:295–300

Zhang Y, Rózsa M, Liang Y, Bushey D, Wei Z, Zheng J, Reep D, Broussard GJ, Tsang A, Tsegaye G, et al. (2023) Fast and sensitive GCaMP calcium indicators for imaging neural populations. Nature (London).6157954:884–91

Zhang Y, Looger LL. (2024) Fast and sensitive GCaMP calcium indicators for neuronal imaging. The Journal of Physiology.6028:1595–604

Akerboom J, Chen TW, Wardill TJ, Tian L, Marvin JS, Mutlu S, Calderón NC, Esposti F, Borghuis BG, Sun XR, et al. (2012) Optimization of a GCaMP calcium indicator for neural activity imaging. Journal of Neuroscience.3238:13819–40

Peron Simon P, Freeman J, Iyer V, Guo C, Svoboda K. (2015) A Cellular Resolution Map of Barrel Cortex Activity during Tactile Behavior. Neuron.863:783–99

Ziv Y, Burns LD, Cocker ED, Hamel EO, Ghosh KK, Kitch LJ, Gamal AE, Schnitzer MJ. (2013) Long-term dynamics of CA1 hippocampal place codes. Nature Neuroscience.163:264–6

McMahon SM, Jackson MB. (2018) An Inconvenient Truth: Calcium Sensors Are Calcium Buffers. Trends Neurosci.4112:880–4

Zhang D, Redington E, Gong Y. (2021) Rational engineering of ratiometric calcium sensors with bright green and red fluorescent proteins. Communications Biology.41:924

Bellen HJ, Tong C, Tsuda H. (2010) 100 years of Drosophila research and its impact on vertebrate neuroscience: a history lesson for the future. Nat Rev Neurosci.117:514–22

Frank CA, Wang X, Collins CA, Rodal AA, Yuan Q, Verstreken P, Dickman DK. (2013) New approaches for studying synaptic development, function, and plasticity using Drosophila as a model system. J Neurosci.3345:17560–8

Macleod GT, Hegström-Wojtowicz M, Charlton MP, Atwood HL. (2002) Fast Calcium Signals in Drosophila Motor Neuron Terminals. Journal of Neurophysiology.885:2659–63

Kiragasi B, Wondolowski J, Li Y, Dickman DK. (2017) A Presynaptic Glutamate Receptor Subunit Confers Robustness to Neurotransmission and Homeostatic Potentiation. Cell Rep.1913:2694–706

Cohn R, Morantte I, Ruta V. (2015) Coordinated and Compartmentalized Neuromodulation Shapes Sensory Processing in Drosophila. Cell.1637:1742–55

Akbergenova Y, Cunningham KL, Zhang YV, Weiss S, Littleton JT. (2018) Characterization of developmental and molecular factors underlying release heterogeneity at Drosophila synapses. eLife.7e38268

Newman ZL, Hoagland A, Aghi K, Worden K, Levy SL, Son JH, Lee LP, Isacoff EY. (2017) Input-Specific Plasticity and Homeostasis at the Drosophila Larval Neuromuscular Junction. Neuron.936:1388–404.e10

Peled ES, Newman ZL, Isacoff EY. (2014) Evoked and spontaneous transmission favored by distinct sets of synapses. Curr Biol.245:484–93

Perry S, Han Y, Qiu C, Chien C, Goel P, Nishimura S, Sajnani M, Schmid A, Sigrist SJ, Dickman D. (2022) A glutamate receptor C-tail recruits CaMKII to suppress retrograde homeostatic signaling. Nature Communications.131:

Li X, Chien C, Han Y, Sun Z, Chen X, Dickman D. (2021) Autocrine inhibition by a glutamate-gated chloride channel mediates presynaptic homeostatic depression. Science advances.749:eabj1215-eabj

Han Y, Goel P, Chen J, Perry S, Tran N, Nishimura S, Sanjani M, Chien C, Dickman D. (2023) Excess glutamate release triggers subunit-specific homeostatic receptor scaling. Cell Reports.427:112775

Bindels DS, Haarbosch L, van Weeren L, Postma M, Wiese KE, Mastop M, Aumonier S, Gotthard G, Royant A, Hink MA, et al. (2017) mScarlet: a bright monomeric red fluorescent protein for cellular imaging. Nature Methods.141:53–6

Gadella TWJ, van Weeren L, Stouthamer J, Hink MA, Wolters AHG, Giepmans BNG, Aumonier S, Dupuy J, Royant A. (2023) mScarlet3: a brilliant and fast-maturing red fluorescent protein. Nature Methods.204:541–5

Schmid A, Hallermann S, Kittel RJ, Khorramshahi O, Frölich AM, Quentin C, Rasse TM, Mertel S, Heckmann M, Sigrist SJ. (2008) Activity-dependent site-specific changes of glutamate receptor composition in vivo. Nat Neurosci.116:659–66

Han Y, Chien C, Goel P, He K, Pinales C, Buser C, Dickman D. (2022) Botulinum neurotoxin accurately separates tonic vs. phasic transmission and reveals heterosynaptic plasticity rules in Drosophila. eLife.11

Müller M, Davis GW. (2012) Transsynaptic control of presynaptic Ca²⁺ influx achieves homeostatic potentiation of neurotransmitter release. Curr Biol.2212:1102–8

Giovannucci A, Friedrich J, Gunn P, Kalfon J, Brown BL, Koay SA, Taxidis J, Najafi F, Gauthier JL, Zhou P, et al. (2019) CaImAn an open source tool for scalable calcium imaging data analysis. eLife.8e38173

Blum ID, Keleş MF, Baz E-S, Han E, Park K, Luu S, Issa H, Brown M, Ho MCW, Tabuchi M, et al. (2021) Astroglial Calcium Signaling Encodes Sleep Need in Drosophila. Current Biology.311:150–62.e7

Xing X, Wu C-F. (2018) Unraveling Synaptic GCaMP Signals: Differential Excitability and Clearance Mechanisms Underlying Distinct Ca^2+^ Dynamics in Tonic and Phasic Excitatory, and Aminergic Modulatory Motor Terminals in *Drosophila*. eneuro.51:ENEURO.0362-17.2018

Reagents G, Hasseman JP, Turner GC, Looger LL, Bushey D, Reep D, Tsang A, Zheng J, Korff W, Jayaraman V, et al. jGCaMP8 Variants with Improved Expression for Drosophila 2024. Available from: 10.6084/m9.figshare.26231087.v1.

Lu Z, Chouhan AK, Borycz JA, Lu Z, Rossano AJ, Brain KL, Zhou Y, Meinertzhagen IA, Macleod GT. (2016) High-Probability Neurotransmitter Release Sites Represent an Energy-Efficient Design. Curr Biol.2619:2562–71

He K, Han Y, Li X, Hernandez RX, Riboul DV, Feghhi T, Justs KA, Mahneva O, Perry S, Macleod GT, et al. (2023) Physiologic and Nanoscale Distinctions Define Glutamatergic Synapses in Tonic vs Phasic Neurons. The Journal of neuroscience.4325:4598–611

He K, Dickman D. (2025) Building and modifying diverse synaptic properties: Insights from Drosophila. Curr Opin Neurobiol.92102995

Davis GW, Müller M. (2015) Homeostatic control of presynaptic neurotransmitter release. Annu Rev Physiol.77251–70

Goel P, Dickman D. (2021) Synaptic homeostats: latent plasticity revealed at the Drosophila neuromuscular junction. Cell Mol Life Sci.787:3159–79

Delvendahl I, Müller M. (2019) Homeostatic plasticity-a presynaptic perspective. Curr Opin Neurobiol.54155–62

Chien C, He K, Perry S, Tchitchkan E, Han Y, Li X, Dickman D. (2025) Distinct input-specific mechanisms enable presynaptic homeostatic plasticity. Sci Adv.117:eadr0262

Maravall M, Mainen ZF, Sabatini BL, Svoboda K. (2000) Estimating intracellular calcium concentrations and buffering without wavelength ratioing. Biophys J.785:2655–67

Sinha SR, Wu LG, Saggau P. (1997) Presynaptic calcium dynamics and transmitter release evoked by single action potentials at mammalian central synapses. Biophys J.722 Pt 1:637–51

Helmchen F, Borst JG, Sakmann B. (1997) Calcium dynamics associated with a single action potential in a CNS presynaptic terminal. Biophys J.723:1458–71

Ramesh N, Escher MJF, Mampell MM, Böhme MA, Götz TWB, Goel P, Matkovic T, Petzoldt AG, Dickman D, Sigrist SJ. (2021) Antagonistic interactions between two Neuroligins coordinate pre- and postsynaptic assembly. Curr Biol.318:1711–25.e5

Chen J, He K, Han Y, Dickman D. (2024) Ca(2+) imaging of synaptic compartments using subcellularly targeted GCaMP8f in Drosophila. STAR Protoc.51:102832

Grienberger C, Konnerth A. (2012) Imaging Calcium in Neurons. Neuron.735:862–85

Hendel T, Mank M, Schnell B, Griesbeck O, Borst A, Reiff DF. (2008) Fluorescence changes of genetic calcium indicators and OGB-1 correlated with neural activity and calcium in vivo and in vitro. J Neurosci.2829:7399–411

Lock JT, Parker I, Smith IF. (2015) A comparison of fluorescent Ca²⁺ indicators for imaging local Ca²⁺ signals in cultured cells. Cell Calcium.586:638–48

Macleod GT. (2012) Imaging and analysis of single synapse Ca²⁺ dynamics in the larval Drosophila neuromuscular junction. Cold Spring Harbor Protocols.201212:1277–83

Lnenicka GA, Grizzaffi J, Lee B, Rumpal N. (2006) Ca^2+^ Dynamics along Identified Synaptic Terminals in *Drosophila* Larvae. The Journal of Neuroscience.2647:12283–93

Justs KA, Lu Z, Chouhan AK, Borycz JA, Lu Z, Meinertzhagen IA, Macleod GT. (2022) Presynaptic Mitochondrial Volume and Packing Density Scale with Presynaptic Power Demand. J Neurosci.426:954–67

Lu C, Li Q, Li Y, Wang Y, Zhang Y-F. (2016) A Short Glance at the Neural Circuitry Mechanism Underlying Depression. World Journal of Neuroscience.63:

Macleod GT. (2012) Imaging and analysis of nonratiometric calcium indicators at the Drosophila larval neuromuscular junction. Cold Spring Harb Protoc.20127:802–9

Cunningham KL, Sauvola CW, Tavana S, Littleton JT. (2022) Regulation of presynaptic Ca2+ channel abundance at active zones through a balance of delivery and turnover. eLife.11e78648

Gratz SJ, Goel P, Bruckner JJ, Hernandez RX, Khateeb K, Macleod GT, Dickman D, O’Connor-Giles KM. (2019) Endogenous Tagging Reveals Differential Regulation of Ca(2+) Channels at Single Active Zones during Presynaptic Homeostatic Potentiation and Depression. J Neurosci.3913:2416–29

Medeiros AT, Gratz SJ, Delgado A, Ritt JT, O’Connor-Giles KM. (2024) Ca2+ channel and active zone protein abundance intersects with input-specific synapse organization to shape functional synaptic diversity. eLife.12RP88412

Kikuma K, Li X, Perry S, Li Q, Goel P, Chen C, Kim D, Stavropoulos N, Dickman D. (2019) Cul3 and insomniac are required for rapid ubiquitination of postsynaptic targets and retrograde homeostatic signaling. Nat Commun.101:2998

Perry S, Han Y, Das A, Dickman D. (2017) Homeostatic plasticity can be induced and expressed to restore synaptic strength at neuromuscular junctions undergoing ALS-related degeneration. Hum Mol Genet.2621:4153–67

Virtanen P, Gommers R, Oliphant TE, Haberland M, Reddy T, Cournapeau D, Burovski E, Peterson P, Weckesser W, Bright J, et al. (2020) SciPy 1.0: fundamental algorithms for scientific computing in Python. Nature Methods.173:261–72

Shemesh OA, Linghu C, Piatkevich KD, Goodwin D, Celiker OT, Gritton HJ, Romano MF, Gao R, Yu CJ, Tseng HA, et al. (2020) Precision Calcium Imaging of Dense Neural Populations via a Cell-Body-Targeted Calcium Indicator. Neuron.1073:470–86.e11

Fleming W, Jewell S, Engelhard B, Witten DM, Witten IB. (2021) Inferring spikes from calcium imaging in dopamine neurons. PLoS One.166:e0252345

