## Supplemental Table 1 for "Resolving synaptic events using subsynaptically targeted GCaMP8 variants"

**Supplementary Table 1: Absolute values and statistical comparisons for $Ca^{2+}$ imaging and electrophysiology data.**
This table reports the full statistical details and properties for data presented in the indicated figures, including p-values, mean ± SEM, sample sizes (n), and genotypes for all conditions tested. $Ca^{2+}$ imaging parameters include ΔF/F (or ΔR/R), rise time ($\tau_{rise}$), and decay time ($\tau_{decay}$) constants. Electrophysiological parameters include mEPSP amplitude, EPSP amplitude, quantal content (QC), input resistance, and resting potential. p-values from one-way ANOVA with Tukey's multiple comparison test are shown for key contrasts between genotypes and indicators. Data for outlier analysis, correlation analysis and linear regression analyses are included where applicable.

| Figure | Label | Genotype | Motor Neuron | △F/F (△R/R) | p value | Tau rise (msec) | p value | Tau decay (msec) | p value | n |
|---|---|---|---|---|---|---|---|---|---|---|
| 3B | OGB-1 | | Ib | 0.227 (±0.020) | 0.0279 (*) | 3.040 (±0.291) | 0.7398 (ns) | 60.13 (±3.259) | <0.0001 (****) | 7 |
| 3B | Scar8f | *w;OK6-GAL4/+; UAS-Syt::mScarlet::GCaMP8f/+* | Ib | 0.322 (±0.032) | | 2.893 (±0.323) | | 38.52 (±1.053) | | 7 |
| 3D | RSET-GCaMP8m | *w;OK319-GAL4/+; UAS-RSET-GCaMP8m/+* | Ib | 0.397 (±0.029) | <0.0001 (****) | 6.933 (±0.454) | >0.9999 (ns) | 137.8 (±6.887) | 0.2679 (ns) | 7 |
| 3D | Scar8m | *w;OK319-GAL4/+; UAS-Syt::mScarlet3::GCaMP8m/+* | Ib | 0.6765 (±0.018) | | 6.736 (±0.452) | | 100.8 (±5.135) | | 13 |

| Figure | Label | Genotype | Motor Neuron | △F/F (△R/R) | p value (6s vs 8f, 6s vs 8m, 8f vs 8m) | Tau rise (msec) | p value (6s vs 8f, 6s vs 8m, 8f vs 8m) | Tau decay (msec) | p value (6s vs 8f, 6s vs 8m, 8f vs 8m) | n |
|---|---|---|---|---|---|---|---|---|---|---|
| 4E | Syt::GCaMP6s | *w;OK319-GAL4/+; UAS-Syt::GCaMP6s/+* | Ib | 0.138 (±0.004) | <0.0001 (****), <0.0001 (****), <0.0001 (****) | 43.77 (±3.134) | <0.0001 (****), <0.0001 (****), 0.7827 (ns) | 171.7 (±8.01) | <0.0001 (****), <0.0001 (****), 0.0046 (**) | 14 |
| 4E | Scar8f | *w;OK319-GAL4/+; UAS-Syt::mScarlet::GCaMP8f/+* | Ib | 0.389 (±0.010) | | 5.265 (±1.165) | | 66.64 (±1.849) | | 15 |
| 4E | Scar8m | *w;OK319-GAL4/+; UAS-Syt:: mScarlet3::GCaMP8m/+* | Ib | 0.632 (±0.011) | | 6.984 (±0.604) | | 99.20 (±3.785) | | 14 |

| Figure | Label | Genotype | Motor Neuron | △F/F (△R/R) | p value | n |
|---|---|---|---|---|---|---|
| 5D | Scar8m/WT Ib | *w;OK319-GAL4/+; UAS-Syt:: mScarlet3::GCaMP8m/+* | Ib | 0.590 (±0.048) | <0.0001 (****) | 9 |
| 5D | Scar8m/WT Is | *w;OK319-GAL4/+; UAS-Syt:: mScarlet3::GCaMP8m/+* | Is | 1.100 (±0.060) | | 9 |
| 5F | Scar8m/WT Ib | w;OK319-GAL4/+; UAS-Syt:: mScarlet3::GCaMP8m/+ | Ib | 0.605 (±0.042) | <0.0001 (****) | 9 |
| 5F | GluRIIA$^{-/-}$ Ib | *W;OK371,GluRIIA$^{pv3}$/GluRIIA$^{pv3}$; UAS-Syt:: mScarlet3::GCaMP8m/+* | Ib | 0.941 (±0.035) | | 12 |

| Figure | Label | Genotype | Motor Neuron | △F/F (△R/R) | p value (8f vs 8m area, 8f vs 8m line, 8m line vs 8m area) | Tau rise (msec) | p value (8f vs 8m area, 8f vs 8m line, 8m line vs 8m area) | Tau decay (msec) | p value (8f vs 8m area, 8f vs 8m line, 8m line vs 8m area) | n |
|---|---|---|---|---|---|---|---|---|---|---|
| 6D | Bar8f | *w;OK6-GAL4/+; BRP::mScarlet::GCaMP8f/+* | Ib | 1.904 (±0.298) | <0.0001 | 9.396 (±2.151) | 0.822 (ns), | 22.71 (±18.553) | <0.0001 | 8 |

| | | | | | | | | | | |
|---|---|---|---|---|---|---|---|---|---|---|
| 6D | Bar8m (area scan) | *w;OK6-GAL4/+; BRP::mScarlet::GCaMP8m/+* | Ib | 3.280 (±0.415) | (****), 0.357, (ns), <0.0001 (****) | 9.019 (±2.575) | <0.0001 (****), <0.0001 (****) | 47.97 (±10.973) | (****), <0.0001 (****), 0.134 (ns) | 7 |
| 6D | Bar8m (line scan) | *w;OK6-GAL4/+; BRP::mScarlet::GCaMP8m/+* | Ib | 2.051 (±0.503) | | 3.230 (±1.471) | | 40.26 (±9.812) | | 8 |

**Figure 6E (left):** Data represent mean ± SEM of ΔR/R values for Resonant area scan of 30 active zones from Bar8f-expressing NMJs (genotype: *w;OK6-GAL4/+; BRP::mScarlet::GCaMP8f/+*) (n=8 NMJ). Outlier status is determined using the interquartile range (IQR) method on individual data points: Q1 = 1.454 (25th percentile), Q3 = 2.268 (75th percentile). Mean intensities of columns 27–30 exceed the upper bound and are marked as outliers.

| | | | | | | | | | | |
|---|---|---|---|---|---|---|---|---|---|---|
| **AZ#** | **1** | **2** | **3** | **4** | **5** | **6** | **7** | **8** | **9** | **10** |
| **△R/R** | 1.504 | 1.513 | 1.549 | 1.566 | 1.644 | 1.661 | 1.691 | 1.697 | 1.701 | 1.720 |
| **Std. Deviation** | 0.07174 | 0.1498 | 0.1143 | 0.1017 | 0.1038 | 0.08544 | 0.1460 | 0.08549 | 0.09921 | 0.1067 |
| **Outlier** | No | No | No | No | No | No | No | No | No | No |
| **AZ#** | 10 | 12 | 13 | 14 | 15 | 16 | 17 | 18 | 19 | 20 |
| **△R/R** | 1.755 | 1.759 | 1.785 | 1.787 | 1.807 | 1.824 | 1.896 | 1.900 | 1.907 | 1.925 |
| **Std. Deviation** | 0.1229 | 0.1049 | 0.1203 | 0.1668 | 0.1528 | 0.06702 | 0.1379 | 0.1195 | 0.1791 | 0.1929 |
| **Outlier** | No | No | No | No | No | No | No | No | No | No |
| **AZ#** | 21 | 22 | 23 | 24 | 25 | 26 | 27 | 28 | 29 | 30 |
| **△R/R** | 2.023 | 2.062 | 2.072 | 2.127 | 2.134 | 2.184 | 2.377 | 2.413 | 2.425 | 2.706 |
| **Std. Deviation** | 0.2157 | 0.1229 | 0.1357 | 0.1389 | 0.1387 | 0.1848 | 0.2230 | 0.1628 | 0.1804 | 0.1372 |
| **Outlier** | No | No | No | No | No | No | Yes | Yes | Yes | Yes |

**Additional info for Figure 6E: Additional analysis of calcium signal amplitudes versus AZ size for Bar8f.** (Left) Scatter plots relating AZ size (measured by mScarlet1 fluorescence area) to ΔR/R. A weak negative correlation is observed between ΔR/R and AZ size, likely due to overestimation of ROI boundaries during manual AZ segmentation. (Right) Relationship between active zone (AZ) size and peak ΔR/R responses measured with Bar8f in Ib motor neurons. Each point represents an individual AZ (n = 29). A modest negative correlation was observed, likely due to overestimation of ROI boundaries during manual AZ segmentation. The best-fit linear regression is shown in red.

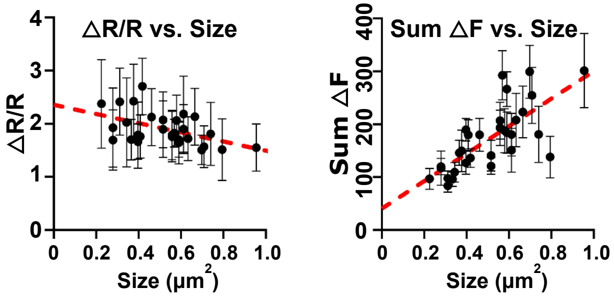

| Figure | Label | Genotype | Motor Neuron | Metric (Y vs X) | Equation | Pearson's r | R² | p-value (correlation) | n |
|---|---|---|---|---|---|---|---|---|---|
| Additional info for Figure 6 | Bar8f | *w;OK6-GAL4/+; BRP::mScarlet::GCaMP8f/+* | Ib | ΔR/R vs. AZ Size | Y = -0.8644*X + 2.355 | -0.4902 | 0.2403 | 0.0069 (**) | 29 |
| | Bar8f | *w;OK6-GAL4/+; BRP::mScarlet::GCaMP8f/+* | Ib | Sum ΔF vs. AZ Size | Y = 259.8*X + 40.40 | 0.7316 | 0.5353 | <0.0001 (****) | 29 |

**Figure 6E (middle):** Data represent mean ± SEM of ΔR/R values for Resonant area scan of 30 active zones from Bar8m-expressing NMJs (genotype: *w;OK6-GAL4/+; BRP::mScarlet::GCaMP8m/+*) (n=7 NMJ). Outlier status is determined using the interquartile range (IQR) method on individual data points: Q1 = 2.71 (25th percentile), Q3 = 3.83 (75th percentile). The mean intensity values in columns 1, 28, 29, and 30 exceed the upper bound and are marked as outliers.

| AZ# | 1 | 2 | 3 | 4 | 5 | 6 | 7 | 8 | 9 | 10 |
|---|---|---|---|---|---|---|---|---|---|---|
| △R/R | 2.570 | 2.711 | 2.718 | 2.777 | 2.908 | 2.920 | 2.962 | 3.006 | 3.008 | 3.011 |
| Std. Deviation | 0.1135 | 0.1574 | 0.09472 | 0.4138 | 0.08930 | 0.1761 | 0.1111 | 0.09609 | 0.1770 | 0.3050 |
| Outlier | Yes | No | No | No | No | No | No | No | No | No |
| AZ# | 10 | 12 | 13 | 14 | 15 | 16 | 17 | 18 | 19 | 20 |
| △R/R | 3.014 | 3.023 | 3.055 | 3.222 | 3.275 | 3.312 | 3.338 | 3.344 | 3.372 | 3.423 |
| Std. Deviation | 0.2792 | 0.2036 | 0.2087 | 0.2315 | 0.1654 | 0.2702 | 0.1639 | 0.1447 | 0.1348 | 0.2328 |
| Outlier | No | No | No | No | No | No | No | No | No | No |
| AZ# | 21 | 22 | 23 | 24 | 25 | 26 | 27 | 28 | 29 | 30 |
| △R/R | 3.438 | 3.481 | 3.529 | 3.587 | 3.624 | 3.692 | 3.831 | 3.935 | 4.146 | 4.160 |
| Std. Deviation | 0.2824 | 0.2898 | 0.2229 | 0.2349 | 0.2161 | 0.2328 | 0.1887 | 0.2629 | 0.2488 | 0.2867 |
| Outlier | No | No | No | No | No | No | No | Yes | Yes | Yes |

**Figure 6E (right):** Data represent mean ± SEM of ΔR/R values for Galvano line scan of 30 active zones from Bar8m-expressing NMJs (genotype: *w;OK6-GAL4/+; BRP::mScarlet::GCaMP8m/+*) (n=8 NMJ). Outlier status is determined using the interquartile range (IQR) method on individual data points: Q1 = 1.65 (25th percentile), Q3 = 2.46 (75th percentile). The mean intensity values in column 1, 28, 29, 30 exceeds the upper bound and is marked as outliers.

| AZ# | 1 | 2 | 3 | 4 | 5 | 6 | 7 | 8 | 9 | 10 |
|---|---|---|---|---|---|---|---|---|---|---|
| △R/R | 1.137 | 1.377 | 1.584 | 1.627 | 1.642 | 1.642 | 1.657 | 1.677 | 1.709 | 1.752 |
| Std. Deviation | 0.02965 | 0.02603 | 0.04381 | 0.05424 | 0.02733 | 0.07045 | 0.04215 | 0.04927 | 0.04375 | 0.04886 |
| Outlier | Yes | Yes | Yes | Yes | Yes | Yes | No | No | No | No |
| AZ# | 10 | 12 | 13 | 14 | 15 | 16 | 17 | 18 | 19 | 20 |
| △R/R | 1.782 | 1.803 | 1.803 | 1.814 | 1.865 | 1.917 | 1.958 | 2.107 | 2.114 | 2.127 |
| Std. Deviation | 0.04090 | 0.04185 | 0.04185 | 0.03195 | 0.04062 | 0.05031 | 0.06363 | 0.1014 | 0.09142 | 0.07513 |

| | | | | | | | | | | |
|---|---|---|---|---|---|---|---|---|---|---|
| **Outlier** | No | No | No | No | No | No | No | No | No | No |
| **AZ#** | 21 | 22 | 23 | 24 | 25 | 26 | 27 | 28 | 29 | 30 |
| **△R/R** | 2.210 | 2.265 | 2.320 | 2.455 | 2.539 | 2.712 | 2.914 | 2.939 | 3.027 | 3.064 |
| **Std. Deviation** | 0.09016 | 0.07923 | 0.08218 | 0.04884 | 0.08143 | 0.07516 | 0.1091 | 0.09999 | 0.07484 | 0.1087 |
| **Outlier** | No | No | No | No | Yes | Yes | Yes | Yes | Yes | Yes |

| Figure | Label | Genotype | Motor Neuron | △F/F | p value (6s vs 8f, 6s vs 8m, 8f vs 8m) | Tau rise (msec) | p value (6s vs 8f, 6s vs 8m, 8f vs 8m) | Tau decay (msec) | p value (6s vs 8f, 6s vs 8m, 8f vs 8m) | n |
|---|---|---|---|---|---|---|---|---|---|---|
| 7E | SynapGCaMP6f | *w;MHC-CD8-GCaMP6f-Sh;+* | Ib | 0.271 (±0.024) | 0.2164 (ns), <0.0001 (****), <0.0001 (****) | 21.09 (±2.514) | 0.0182 (*), 0.0070 (**), 0.9944 (ns) | 98.76 (±6.914) | <0.0001 (****), <0.0001 (****), 0.0036 (**) | 47 |
| 7E | SynapGCaMP8f | *w;;MHC-CD8-GCaMP8f-Sh* | Ib | 0.349 (±0.030) | | 14.04 (±1.190) | | 41.98 (±1.829) | | 50 |
| 7E | SynapGCaMP8m | *w;;MHC-CD8-GCaMP8m-Sh* | Ib | 0.575 (±0.043) | | 14.22 (±0.985) | | 67.35 (±1.640) | | 58 |

| Figure | Label | Genotype | Motor Neuron | % mEPSPs detected as $Ca^{2+}$ minis | p value (6s vs 8f, 6s vs 8m, 8f vs 8m) | n |
|---|---|---|---|---|---|---|
| 8B | SynapGCaMP6f | *w;MHC-CD8-GCaMP6f-Sh;+* | Ib | 56.87 (±7.217) | <0.0001 (****), <0.0001 (****), 0.5645 (ns) | 12 |
| 8B | SynapGCaMP8f | *w;;MHC-CD8-GCaMP8f-Sh* | Ib | 87.55 (±1.538) | | 17 |
| 8B | SynapGCaMP8m | *w;;MHC-CD8-GCaMP8m-Sh* | Ib | 93.08 (±1.318) | | 12 |

| Figure | Label | Genotype | Motor Neuron | Metric (Y vs X) | Equation | Pearson's r | $R^2$ | p-value (correlation) | n |
|---|---|---|---|---|---|---|---|---|---|
| 8C | SynapGCaMP6f | *w;MHC-CD8-GCaMP6f-Sh;+* | Ib | $Ca^{2+}$ mini ΔF/F vs. mEPSP (mV) | Y = 0.1231*X + 0.08230 | 0.4579 | 0.2097 | <0.0001 (****) | 83 |
| 8C | SynapGCaMP8f | *w;;MHC-CD8-GCaMP8f-Sh* | Ib | $Ca^{2+}$ mini ΔF/F vs. mEPSP (mV) | Y = 0.2910*X + 0.1663 | 0.7316 | 0.5374 | <0.0001 (****) | 98 |
| 8D | SynapGCaMP8m | *w;;MHC-CD8-GCaMP8m-Sh* | Ib | $Ca^{2+}$ mini ΔF/F vs. mEPSP (mV) | Y = 0.4216*X + 0.1672 | 0.8098 | 0.6557 | <0.0001 (****) | 97 |

| Figure | Label | Genotype | Motor Neuron | mEPSP amplitude (mV) | mEPSP frequency (Hz) | R input (MΩ) | Resting potential (mV) | P Value (significance: mEPSP, mEPSP freq) | n |
|---|---|---|---|---|---|---|---|---|---|
| 8E | WT | *w;;Is-GAL4/UAS-BoNT-C* | Ib | 0.636 (±0.015) | 1.062 (±0.100) | 11.91 (±0.115) | 64.45 (±1.561) | - | 41 |
| 8E | $IIB^{-/-}$ | *w;GluRIIB*$^{sp5}$*;Is-GAL4/UAS-BoNT-C* | Ib | 0.824 (±0.017) | 2.433 (±0.147) | 12.89 (±0.197) | 65.46 (±1.541) | <0.0001 (****), <0.0001 (****) | 41 |
| 8E | $IIA^{-/-}$ | *w;GluRIIA*$^{pv3}$*;Is-GAL4/UAS-BoNT-C* | Ib | 0.383 (±0.007) | 0.715 (±0.111) | 12.11 (±0.138) | 65.38 (±1.19) | <0.0001 (****), <0.0001 (****) | 41 |

| Figure | Label | Genotype | Motor Neuron | △F/F | p value (WT vs IIB-/-, WT vs IIA-/-, IIB-/- vs IIA-/-) | n |
|---|---|---|---|---|---|---|
| 8E | WT | *w;;MHC-CD8-GCaMP8m-Sh* | Ib | 0.713 (±0.021) | <0.0001 (****), <0.0001 (****), <0.0001 (****) | 65 |
| 8E | $IIB^{-/-}$ | *W;GluRIIBsp5;MHC-CD8-GCaMP8m-Sh* | Ib | 0.967 (±0.036) | | 69 |
| 8E | $IIA^{-/-}$ | *W;GluRIIBsp5;MHC-CD8-GCaMP8m-Sh* | Ib | 0.427 (±0.017) | | 59 |

**Kolmogorov–Smirnov Test Results for Mini Amplitude Distributions for MN-Ib**

D statistic: Maximum difference between cumulative distributions.

All comparisons reached $p < 0.001$, indicating significantly different mini amplitude distributions.

| Figure | Label | Genotype | Motor Neuron | mEPSP amplitude (mV) | D statistic (WT vs IIB-/-, WT vs IIA-/-, IIB-/- vs IIA-/-) | P Value (WT vs IIB-/-, WT vs IIA-/-, IIB-/- vs IIA-/-) | n |
|---|---|---|---|---|---|---|---|
| 8F | WT | *w;;Is-GAL4/UAS-BoNT-C* | Ib | 0.636 (±0.015) | 0.1854, 0.3647, 0.4043 | <0.0001 (****), <0.0001 (****), <0.0001 (****) | 41 |
| 8F | $IIB^{-/-}$ | *w;GluRIIB*$^{sp5}$*;Is-GAL4/UAS-BoNT-C* | Ib | 0.824 (±0.017) | | | 41 |
| 8F | $IIA^{-/-}$ | *w;GluRIIA*$^{pv3}$*;Is-GAL4/UAS-BoNT-C* | Ib | 0.383 (±0.007) | | | 41 |

| Figure | Label | Genotype | Motor Neuron | △F/F | D statistic (WT vs IIB-/-, WT vs IIA-/-, IIB-/- vs IIA-/-) | p value (WT vs IIB-/-, WT vs IIA-/-, IIB-/- vs IIA-/-) | n |
|---|---|---|---|---|---|---|---|
| 8F | WT | *w;;MHC-CD8-GCaMP8m-Sh* | Ib | 0.713 (±0.021) | 0.2046, 0.2938, 0.3277 | <0.0001 (****), <0.0001 (****), <0.0001 (****) | 65 |
| 8F | $IIB^{-/-}$ | *W;GluRIIB*$^{sp5}$*;MHC-CD8-GCaMP8m-Sh* | Ib | 0.967 (±0.036) | | | 69 |
| 8F | $IIA^{-/-}$ | *W;GluRIIB*$^{sp5}$*;MHC-CD8-GCaMP8m-Sh* | Ib | 0.427 (±0.017) | | | 59 |

| Figure | Label | Genotype | Motor Neuron | mEPSP amplitude (mV) | EPSP amplitude (mV) | QC | mEPSP frequency (Hz) | R input (MΩ) | Resting potential (mV) | n | P value (significance: mEPSP, EPSP, QC) |
|---|---|---|---|---|---|---|---|---|---|---|---|
| S1B | WT | *w*$^{1118}$ | Is + Ib | 0.995 (±0.048) | 30.96 (±1.182) | 30.34 (±2.101) | 3.133 (±0.319) | 12.11 (±0.138) | 65.38 (±1.19) | 10 | - |
| S1B | OK319> Syt::GCaMP6s | *w;OK319-GAL4/+; UAS-Syt::GCaMP6s/+* | Is + Ib | 0.996 (±0.057) | 29.39 (±1.035) | 29.88 (±1.024) | 4.067 (±0.509) | 12.89 (±0.197) | 65.46 (±1.541) | 8 | >0.9999 (ns), 0.8985 (ns), 0.9997(ns), 0.9694 (ns) |
| S1B | OK319> Scar8f | *w;OK319-GAL4/+; UAS-Syt::mScarlet::GCaMP8f/+* | Is + Ib | 1.057 (±0.053) | 22.33 (±0.941) | 21.02 (±1.488) | 2.222 (±0.4923) | 12.71 (±0.126) | 66.62 (±1.284) | 9 | 0.7858 (ns), <0.0001 (****), <0.0001 (***), 0.8299 (ns) |
| S1B | OK319> Scar8m | *w;OK319-GAL4/+; UAS-Syt::mScarlet3::GCaMP8m/+* | Is + Ib | 1.031 (±0.027) | 28.61 (±2.122) | 27.89 (±2.289) | 2.692 (±0.187) | 10.87 (±0.108) | 66.23 (±1.105) | 8 | 0.9867 (ns), 0.5656 (ns), 0.7704 (ns), 0.9940 (ns) |
| S1B | OK6> Bar8f | *w;OK6-GAL4/+; BRP::mScarlet::GCaMP8f/+* | Is + Ib | 1.088 (±0.018) | 32.30 (±0.425) | 29.73 (±0.595) | 2.117 (±0.611) | 11.25 (±0.143) | 63.62 (±1.443) | 8 | 0.3526 (ns), 0.9535 (ns), >0.9999 (ns), 0.6383 (ns) |
| S1B | OK6> Bar8m | *w;OK6-GAL4/+; BRP::mScarlet3::GCaMP8m/+* | Is + Ib | 0.9278 (±0.051) | 31.97 (±0.731) | 35.73 (±2.422) | 2.824 (±0.428) | 13.51 (±0.163) | 66.23 (±1.835) | 8 | 0.7966 (ns), 0.9932 (ns), 0.1136 (ns), 0.9870 (ns) |
| S1B | SynapGCaMP6f | *w;MHC-CD8-GCaMP6f-Sh;+* | Is + Ib | 1.043 (±0.025) | 29.69 (±0.934) | 28.60 (±1.011) | 3.606 (±0.608) | 11.84 (±0.198) | 65.13 (±1.484) | 8 | 0.9317 (ns), 0.9646 (ns), 0.9495 (ns), >0.9999 (ns) |
| S1B | SynapGCaMP8f | *w;;MHC-CD8-GCaMP8f-Sh* | Is + Ib | 1.124 (±0.013) | 29.55 (±0.736) | 26.34 (±0.781) | 2.692 (±0.390) | 13.97 (±0.160) | 66.51 (±1.471) | 8 | 0.0825 (ns), 0.9395 (ns), 0.2607 (ns), 0.9839 (ns) |
| S1B | SynapGCaMP8m | *w;;MHC-CD8-GCaMP8m-Sh* | Is + Ib | 1.007 (±0.024) | 29.73 (±0.901) | 28.60 (±1.011) | 3.587 (±0.546) | 11.90 (±0.105) | 67.39 (±1.276) | 8 | >0.9999 (ns), 0.9702 (ns), 0.9998 (ns), >0.9999 (ns) |
