## Supplemental Table 2 for "Resolving synaptic events using subsynaptically targeted GCaMP8 variants"

### KEY RESOURCES TABLE

| REAGENT/RESOURCE | | SOURCE | IDENTIFIER |
|---|---|---|---|
| **Antibodies** | **Dilution** | | |
| Mouse anti-DLG | 1:50 | Developmental Studies Hybridoma Bank (DSHB) | 4F3 |
| Mouse anti-BRP | 1:100 | DSHB | nc82 |
| Chicken anti-GFP | 1:1000 | Aves Lab | GFP-1010 |
| Rabbit anti-GluRIIC | 1:2000 | (Goel & Dickman, 2018) | |
| Rabbit anti-Syt | 1:2000 | (Mackler et al., 2002) | |
| Alexa Fluor 488 conjugated Donkey anti-chicken secondary antibody | 1:400 | Jackson ImmunoResearch | 703-545-155 |
| Alexa Fluor 488 conjugated Goat anti-Horseradish Peroxidase | 1:400 | Jackson ImmunoResearch | 123-545-021 |
| Alexa Fluor 594 conjugated secondary antibodies | 1:400 | Jackson ImmunoResearch | 715-585-150 |
| Alexa Fluor 647 conjugated Donkey anti-mouse secondary antibody | 1:400 | Jackson ImmunoResearch | 715-605-150 |
| Alexa Fluor 647 conjugated Donkey anti-rabbit secondary antibody | 1:400 | Jackson ImmunoResearch | 711-605-152 |
| Cy3-conjugated Donkey anti-rabbit secondary antibody | 1:400 | Jackson ImmunoResearch | 711-165-152 |
| DyLight 405 conjugated Donkey anti-mouse secondary antibody | 1:400 | Jackson ImmunoResearch | 715-475-150 |
| STAR RED conjugated secondary antibodies | 1:200 | Abberior | STRED-1002 |
| **Chemicals** | **Dilution** | | |
| Oregon Green 488 BAPTA-1 dextran, Potassium Salt, 10,000 MW, Anionic | 5 mM | Thermo Fisher Scientific (Invitrogen, Molecular Probes) | O6798 |
| **Drosophila Strains** | | | |
| UAS-Syt::mScarlet::GCaMP8f (Scar8f) | | (Li et al., 2021) | |
| UAS-Syt::mScarlet3::GCaMP8m (Scar8m) | | (This paper) | |
| UAS-Syt::GCaMP6s | | BDSC | 64414 |
| RSET-GCaMP8m | | BDSC | 605073 |
| SynapGCaMP6f | | (Newman et al., 2017) | |
| SynapGCaMP8f | | (Han et al., 2022) | |
| SynapGCaMP8m | | (This paper) | |
| UAS-BRP::mCherry::GCaMP6s | | (Kiragasi et al., 2017) | |
| UAS-BRP::mScarlet::GCaMP8f (Bar8f) | | (This paper) | |
| UAS-BRP::mScarlet3::GCaMP8m (Bar8m) | | (This paper) | |
| OK6-GAL4 | | (Aberle et al., 2002) | |
| OK319-GAL4 | | (Sweeney et al., 1995) | |
| UAS-BoNT-C | | (Han et al., 2022) | |
| R27E09-GAL4 (Is-Gal4) | | BDSC | 49227 |
| $w^{1118}$ | | BDSC | 5905 |

| | | | |
|---|---|---|---|
| *GluRIIA*$^{pv3}$ | | (Han et al., 2023) | |
| *GluRIIB*$^{sp5}$ | | (Han et al., 2023) | |
| **Software and Algorithms** | | | |
| ImageJ | | https://imagej.net/ | 1.8.0 |
| NIS-Elements software | | Nikon Instruments | 5.41.02 (Build 1711) |
| Huygens Essential | | Scientific Volume Imaging | 25.04 |
| Mini Analysis | | Synaptosoft | 6.0.7 |
| Axon pCLAMP Clampfit | | Molecular Devices | 10.7 |
| Clampex | | Molecular Devices | 10.7 |
| GraphPad Prism | | GraphPad | 10.0.1 |
| Jupyter Notebook | | Anaconda | 6.0.1 |
| Python | | https://www.python.org/ | 3.10.11 |
| Excel | | Microsoft | 2021 |
| CaFire | | (This paper) https://github.com/linj7/CaFire | 2.2.1 |

### REFERENCES


Aberle, H., Haghighi, A. P., Fetter, R. D., McCabe, B. D., Magalhães, T. R., & Goodman, C. S. (2002). wishful thinking encodes a BMP type II receptor that regulates synaptic growth in Drosophila. *Neuron*, *33*(4), 545-558. https://doi.org/10.1016/s0896-6273(02)00589-5

Goel, P., & Dickman, D. (2018). Distinct homeostatic modulations stabilize reduced postsynaptic receptivity in response to presynaptic DLK signaling. *Nature communications*, *9*(1), 1856-1814. https://doi.org/10.1038/s41467-018-04270-0

Han, Y., Chien, C., Goel, P., He, K., Pinales, C., Buser, C., & Dickman, D. (2022). Botulinum neurotoxin accurately separates tonic vs. phasic transmission and reveals heterosynaptic plasticity rules in Drosophila. *eLife*, *11*. https://doi.org/10.7554/eLife.77924

Han, Y., Goel, P., Chen, J., Perry, S., Tran, N., Nishimura, S., Sanjani, M., Chien, C., & Dickman, D. (2023). Excess glutamate release triggers subunit-specific homeostatic receptor scaling. *Cell Reports*.

Kiragasi, B., Wondolowski, J., Li, Y., & Dickman, D. K. (2017). A Presynaptic Glutamate Receptor Subunit Confers Robustness to Neurotransmission and Homeostatic Potentiation. *Cell Rep*, *19*(13), 2694-2706. https://doi.org/10.1016/j.celrep.2017.06.003

Li, X., Chien, C., Han, Y., Sun, Z., Chen, X., & Dickman, D. (2021). Autocrine inhibition by a glutamate-gated chloride channel mediates presynaptic homeostatic depression. *Science advances*, *7*(49), eabj1215-eabj1215. https://doi.org/10.1126/sciadv.abj1215

Mackler, J. M., Drummond, J. A., Loewen, C. A., Robinson, I. M., & Reist, N. E. (2002). The C(2)B Ca(2+)-binding motif of synaptotagmin is required for synaptic transmission in vivo. *Nature*, *418*(6895), 340-344. https://doi.org/10.1038/nature00846

Newman, Z. L., Hoagland, A., Aghi, K., Worden, K., Levy, S. L., Son, J. H., Lee, L. P., & Isacoff, E. Y. (2017). Input-Specific Plasticity and Homeostasis at the Drosophila Larval Neuromuscular Junction. *Neuron*, *93*(6), 1388-1404.e1310. https://doi.org/10.1016/j.neuron.2017.02.028

Sweeney, S. T., Broadie, K., Keane, J., Niemann, H., & O'Kane, C. J. (1995). Targeted expression of tetanus toxin light chain in Drosophila specifically eliminates synaptic transmission and causes behavioral defects. *Neuron*, *14*(2), 341-351. https://doi.org/10.1016/0896-6273(95)90290-2
