## Supplemental Figure 1 for "Resolving synaptic events using subsynaptically targeted GCaMP8 variants"

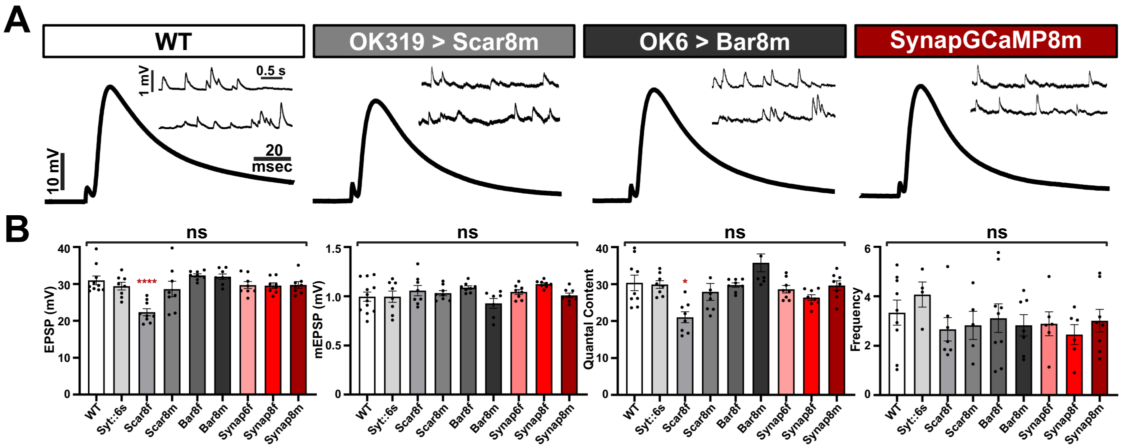


**Supplementary Figure 1: GECI expression does not perturb synaptic transmission at the Drosophila NMJ.**
